# Endocytic recycling is central to circadian collagen fibrillogenesis and disrupted in fibrosis

**DOI:** 10.1101/2021.03.25.436925

**Authors:** Joan Chang, Adam Pickard, Jeremy A. Herrera, Sarah O’Keefe, Richa Garva, John Knox, Thomas A. Jowitt, Matthew Hartshorn, Anna Hoyle, Lewis Dingle, Madeleine Coy, Cédric Zeltz, Jason Wong, Adam Reid, Rajamiyer V. Venkateswaran, Yinhui Lu, Patrick Caswell, Stephen High, Donald Gullberg, Karl E. Kadler

**Author notes:** Co-corresponding authors (JC), (KEK).

## Abstract

Collagen-I fibrillogenesis is crucial to health and development, where dysregulation is a hallmark of fibroproliferative diseases. Here, we show that collagen-I fibril assembly required a functional endocytic system that recycles collagen-I to assemble new fibrils. Endogenous collagen production was not required for fibrillogenesis if exogenous collagen was available, but the circadian-regulated vacuolar protein sorting (VPS) 33b and collagen-binding integrin α11 subunit were crucial to fibrillogenesis. Cells lacking VPS33B secrete soluble collagen-I protomers but were deficient in fibril formation, thus secretion and assembly are separately controlled. Overexpression of VPS33B led to loss of fibril rhythmicity and over-abundance of fibrils, which was mediated through integrin α11β1. Endocytic recycling of collagen-I was enhanced in human fibroblasts isolated from idiopathic pulmonary fibrosis, where VPS33B and integrin α11 subunit were overexpressed at the fibrogenic front; this correlation between VPS33B, integrin α11 subunit, and abnormal collagen deposition was also observed in samples from patients with chronic skin wounds. In conclusion, our study showed that circadian-regulated endocytic recycling is central to homeostatic assembly of collagen fibrils and is disrupted in diseases.

## Introduction

Collagen fibrils account for ~25% of total body protein mass [1] and are the largest protein polymers in vertebrates [2]. The fibrils can exceed centimeters in length and are organized into elaborate networks to provide structural support for cells. It is unclear how the fibrils are formed and how this process goes awry in collagen pathologies, such as fibrosis. A key mechanism, cell surface mediated fibrillogenesis, has been suggested to occur via indirect binding to fibronectin fibrils, or via direct assembly by collagen-binding integrins [3–5]. Interestingly, fibrils can be reconstituted *in vitro* from purified collagen (reviewed in [6]) but the assembly process is not controlled to the extent seen *in vivo* in terms of number, size and organization; this suggests a much tighter cellular control over the process. Additional support for tight cellular control of collagen fibril formation comes from electron microscope observations of collagen fibrils at the plasma membrane of embryonic avian and rodent tendon fibroblasts [7, 8], where the end (tip) of a fibril is enclosed within a plasma membrane invagination termed a fibripositor [9]. Previously we identified vacuolar protein sorting (VPS) 33b (a regulator of SNARE-dependent membrane fusion in the endocytic pathway) as a circadian clock regulated protein involved in collagen homeostasis [10]. VPS33B forms a protein complex with VIPAS39 (VIPAR) [11], and mutations in the *VPS33B* gene cause arthrogryposis-renal dysfunction-cholestasis (ARC) syndrome [12]. Here, death usually occurs within the first year of birth, accompanied with renal insufficiency, jaundice, multiple congenital anomalies, and predisposition to infection [13]. One proposed disease-causing mechanism is abnormal post-Golgi trafficking of lysyl hydroxylase 3 (LH3, PLOD3), which catalyzes the hydroxylation of lysyl residues in collagen to form hydroxylysine residues. It also has hydroxylysyl galactosyltransferase and galactosylhydroxylysyl glucosyltransferase activities, which creates attachment sites for carbohydrates that is crucial in stabilizing intramolecular and intermolecular crosslinks within the collagen structure, and thus is essential for collagen homeostasis during development [14, 15]. All these observations points to a central role for the endosome, situated in proximity of the plasma membrane and acting as a hub for a complex assortment of vesicles, in sorting collagen molecules to different fates.

Previous studies of collagen endocytosis have focused on collagen degradation and signaling, identifying additional collagen-binding proteins such as Endo180 and MRC1 [16–20], as well as a role for non-integrin collagen-receptors involved in mediating signaling activities such as DDR1/DDR2 (reviewed in [21]). Here, we showed that collagen uptake by fibroblasts is circadian rhythmic. Importantly, instead of degrading endocytosed collagen, cells utilize endocytic recycling of exogenous collagen-I to assemble new fibrils, even in the absence of endogenous collagen production. Further we showed that the secretion of soluble collagen protomers is separate from, and can occur independently to, collagen fibril assembly. We identified VPS33B and integrin α11 subunit (part of the collagen-binding integrin α11β1 heterodimer [4, 22, 23]) as central molecules specific to fibril formation. Finally, we showed that in idiopathic pulmonary fibrosis (IPF), a life threatening disease with unknown trigger/mechanism where lung tissue is replaced with collagen fibrils, integrin α11 subunit and VPS33B are located at the invasive fibroblastic focus where collagen fibril rapidly accumulates [24]; IPF fibroblasts also have elevated endocytic recycling of exogenous collagen-I, despite having similar collagen-I expression level to normal fibroblasts. Together, these results provide novel insights into the importance of the circadian clock-regulated endosomal system in normal fibrous tissue homeostasis, where fibrillogenesis occurs with both endogenous collagen and/or recycled scavenged exogenous collagen. This work also highlights that collagen utilization, rather than production (i.e. assembled into fibrils vs. protomeric secretion), is central to the maintenance of homeostasis, and dysregulation leads to disease.

## Results

### Collagen-I is taken up into punctate structures within the cell, and reassembled into fibrils

To study collagen-I endocytosis and the fate of endocytosed collagen, we made Cy3 or Cy5 labeled collagen-I (Cy3-colI, Cy5-colI) from commercial rat-tail collagen [25]. We confirmed the helicity of these collagen with an expected molecular weight corresponding to a heterotrimeric mature collagen-I without the propeptides, and showed that the process of fluorescence-labelling did not alter the collagen trimer secondary structure or stability (**Supplementary Figure 1A-C**). We incubated Cy3-colI with explanted murine tail tendons for 3 days, then imaged the core of the tendon. Cells in tendon (nuclei marked with Hoechst stain) showed a clear uptake of Cy3-colI (**Figure 1A**, yellow box, indicated by yellow arrowheads). Interestingly, fibrillar Cy3 fluorescence signals that extend across cells were also observed, suggestive of these being extracellular fibrils (**Figure 1A**, grey box, indicated by white arrows). Pulse-chase experiments were performed where Cy3-colI was added to the tendons for 3 days, and then removed from the media. This was followed by 5FAM-colI for a further 2 days (**Figure S1D**). The results showed distinct areas where only Cy3-colI was observed (**Figure S1D**, yellow box); this persistence indicated that not all collagen endocytosed by cells is directed for degradation. 5FAM-positive and 5FAM/Cy3-positive fibril-like structures were also observed, confirming that collagen-I taken up by tissues is reincorporated into the matrix (**Figure S1D**, grey boxes). To confirm these findings in 2D cultures, we incubated immortalized murine tendon fibroblast cultures (iTTF) with labelled Cy3-colI. Flow cytometry analysis revealed that most cells had taken up Cy3-colI after overnight incubation (**Figure S1E**). Further time course analyses revealed a time-dependent and concentration-dependent increase of collagen-I uptake (**Figure 1B**). Cells incubated with Cy3-ColI for one hour followed by trypsinization were then analyzed using flow cytometry imaging (flow imaging), revealing that collagen-I is endocytosed by the cells into distinct puncta (**Figure 1C**). Similar results were obtained using Cy5-labelled collagen-I, indicating that the fluorescence label itself did not alter cellular response (**Figure S1F, G, H**). To confirm that the labelled collagen-I is in fact endocytosed into the fibroblasts and not only associated with the cell surface when cells are still attached, we performed live-imaging on cells incubated with Cy3-colI. Time lapse images showed Cy3-colI congregate in structures within the cell, as indicated with white arrows (**Figure S1I**). Additionally, we transduced iTTF with GFP-tagged Rab5 and confirmed co-localisation of Cy5-colI with Rab5-positive intracellular structures (**Figure 1D, yellow arrows**). We then incubated iTTFs with Cy3-colI for one hour, before trypsinizing and replating the cells to ensure any Cy3-colI signal detected hereon originated from the endocytosed pool. Immunofluorescence (IF) staining indicated co-localization of endogenous collagen-I with Cy3-labeled collagen-I in extracellular fibrillar structures after 72hrs, compared to a striking lack of extracellular collagen-I (endogenous or labeled) at 18hrs (**Figure 1E, Figure S2A**). These results demonstrated that exogenous collagen-I can be taken up by cells both *in vitro* and in tissues *ex vivo*, and recycled into fibrils.

**Figure 1:**
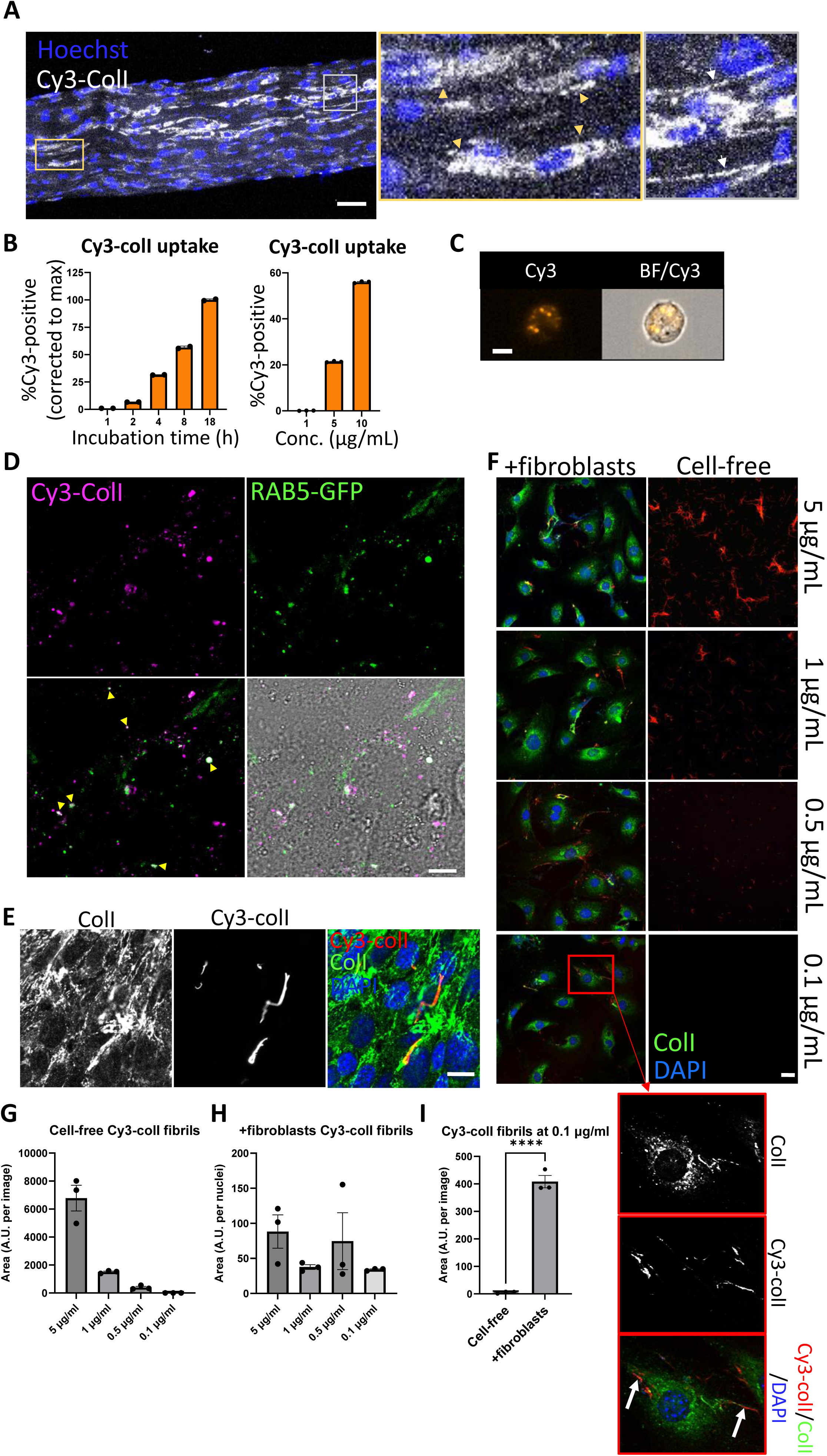
Collagen-I is endocytosed and reassembled into fibrils. A. Fluorescent images of tail tendon incubated with Cy3-colI for 5 days, showing presence of collagen-I within the cells, and fibril-like fluorescence signals outside of cells. Hoechst stain was used to locate cells within the tendon. Area surrounded by yellow box expanded on the right, and cells with Cy3-colI present intracellularly pointed out by yellow triangles. Area surrounded by grey box expanded on the right, and fibril-like fluorescence signals indicated with white arrows. Scale bar = 50 µm. Representative of N = 3. B. Bar chart showing an increase of percentage of fluorescent iTTFs incubated with 1.5 µg/mL Cy3-colI over time (left), and an increase of percentage of fluorescent iTTFs incubated with increasing concentration of Cy3-colI for one hour (right), suggesting a non-linear time-dependent and dose-dependent uptake pattern. N = 3. C. Flow cytometry imaging of iTTFs incubated with 5 µg/mL Cy3-labeled collagen-I for one hour, showing that collagen-I is taken up by cells and held in vesicular-like structures. Images acquired using ImageStream at 40x magnification. Scale bar = 10 *μ*m. Cy3 – Cy3 channel, BF/Cy3 – merged image of BF and Cy3. Representative of >500 cells images collected per condition. D. Fluorescent images of iTTFs transduced with Rab5-GFP and incubated with Cy3-labeled collagen-I. Yellow arrows point to labelled collagen co-localizing with Rab5 in intracellular structures. Representative of N = 3. Scale bar = 10 µm. E. Fluorescent images of iTTFs incubated with 5 µg/mL Cy3-colI for one hour, trypsinized and replated in fresh media, and further cultured for 72 h. Top labels denotes the fluorescence channel corresponding to proteins detected. Merged image colour channels as denoted on top left. Representative of N>3. Scale bar = 20 µm. F. Fluorescent image series of Cy3-colI incubated at different concentrations for 72 h, either cell-free (right panel), or with iTTFs (+fibroblasts, left panel). Representative of N = 3. Scale bar = 20 µm. Red box - zoomed out to the bottom left and separated according to fluorescence channel. White arrows highlighting Cy3-positive fibrils assembled by fibroblasts when incubated with 0.1 µg/mL Cy3-colI. G. Quantification of the area of Cy3-positive fibrils in cell-free cultures, quantified per image area. N=3. H. Quantification of the area of Cy3-positive fibrils in +fibroblasts cultures, corrected to number of nuclei per image area. N=3. I. Comparison of total area of Cy3-positive fibrils in cell-free and +fibroblast cultures at 0.1 µg/mL concentration, as quantified per image area. N=3. *\*\*\*\*p <0.0001*.

We considered the possibility that the fluorescent fibril-like structures were the result of spontaneous cell-free fibrillogenesis of the added Cy3-colI. Thus, we incubated Cy3-colI at concentrations spanning the 0.4 µg/mL critical concentration for cell-free *in vitro* fibril formation [26], with or without cells (**Figure 1F**). In cell-free cultures, Cy3-positive fibrils were observed at 5 µg/mL, 1 µg/mL and 0.5 µg/mL in a dose-dependent manner, with no discernible Cy3-positive fibrils at 0.1 µg/mL Cy3-colI (**Figure 1F**, right panel; **Figure 1G**). However, in the presence of cells, Cy3-positive fibrillar structures were observed at the cell surfaces at all concentrations of Cy3-colI examined, including 0.1 µg/mL (**Figure 1F** zoom ins of red box expanded to the left, highlighted by white arrows; **Figure 1H, I**). These results indicated that the cells actively took up and recycled Cy3-colI into extracellular fibrils.

### The endocytic pathway controls collagen-I secretion and fibril assembly

We then investigated the route that collagen-I is taken up. Prior research identified macropinocytosis [27], which can be modelled with high-molecular weight dextran. We hypothesized that there will be co-localisation of labelled collagen-I and labelled 70kDa dextran within the cell if they are entering through the same route, however surprisingly we see little co-localization (**Figure S2B**). We then performed a receptor saturation experiment, where unlabelled collagen-I was added in excess to labelled collagen-I (10:1) during the 1-hour incubation, before flow imaging analysis of the fibroblasts. Flow imaging showed that labelled collagen was at the cell-periphery and not intracellular when saturation occurs, suggestive of a receptor-mediated process (**Figure S2C**). Thus, we used Dyngo4a [28] to inhibit clathrin-mediated endocytosis, where Dyngo4a treatment (20 µM) leads to ~ 60% reduction in Cy3-colI uptake relative to control (**Figure S2D**), without affecting cell viability (**Figure S2E**). We then investigated the effects of Dyngo4a treatment on the ability of wild-type fibroblasts to assemble collagen fibrils. Cells were treated with Dyngo4a for 48 h before fixation and IF against collagen-I antibody. A significant reduction (average 60%) in the number of collagen-I fibrils assembled was observed (**Figure 2A**), indicative of endocytosis playing a key role in collagen-I fibrillogenesis. Conditioned media (CM) from these treated cells were also collected and assessed. To our surprise, the amount of soluble collagen-I secreted was also greatly reduced (**Figure 2B**). qPCR analyses revealed that *Col1a1* mRNA was significantly reduced in Dyngo4a-treated cells (**Figure S2F**), suggesting a potential feedback mechanism between endocytosis and collagen-I synthesis, secretion, and fibrillogenesis. Interestingly, whilst fibronectin (FN1) mRNA was significantly lower in Dyngo4a-treated cells (**Figure S2F)**, and the intensity of FN1 signal appears lower, the amount of FN1 fibrils deposited (as determined by the area occupied by fibrils) was not significantly impacted (**Figure 2C**). Taken together, these results showed that inhibiting endocytosis in fibroblasts does not lead to accumulation of soluble collagen-I protomers in the extracellular space. Rather, endocytosis impacts collagen fibrillogenesis and the transcriptional control on collagen-I and fibronectin.

**Figure 2:**
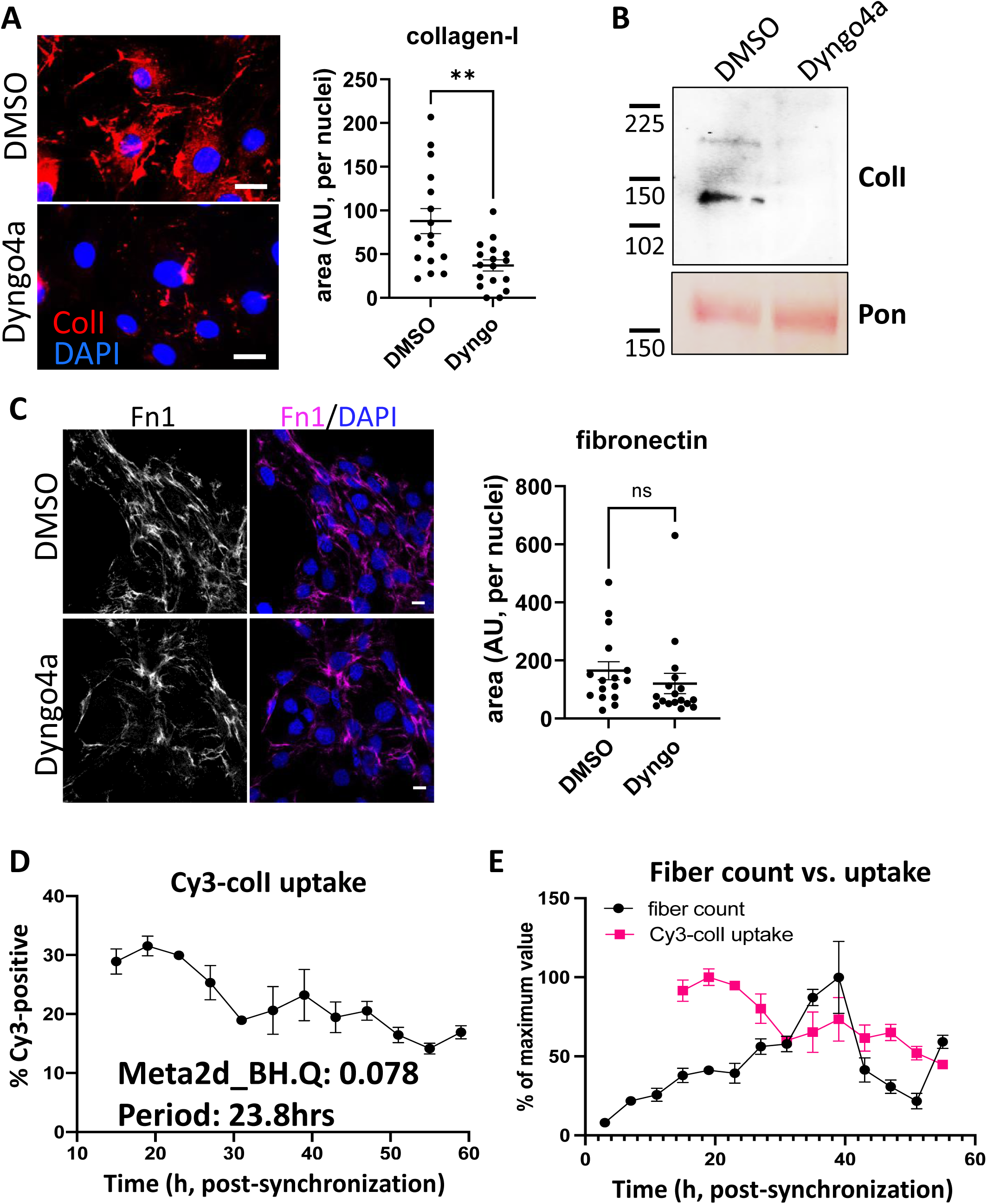
Inhibition of endocytosis leads to changes in collagen-I homeostasis, and endocytosis is a rhythmic event. A. Left: fluorescent images of collagen-I (red) counterstained with DAPI (blue) in iTTFs treated with DMSO (top) or Dyng4a (bottom) for 72 h. Scale bar = 20 µm. Right: quantification of area occupied by collagen-I fibrils, corrected to number of nuclei. N = 3 with 5 images from each experiment ** *p =* 0.0025. B. Western blot analysis of conditioned media taken from iTTFs treated with DMSO or Dyng4a for 72 h, showing a decrease in collagen-I secretion. Top: probed with collagen-I antibody (Col-I), bottom: counterstained with Ponceau (Pon) as control. Protein molecular weight ladders to the left (in kDa). Representative of N = 3. C. Left: fluorescent images of fibronectin (magenta) counterstained with DAPI (blue) in iTTFs treated with DMSO (top) or Dyng4a (bottom) for 72 h. Scale bar = 20 µm. Right: quantification of area occupied by fibronectin fibrils, corrected to number of nuclei. N = 3 with 5 images from each experiment. D. Percentage Cy3-colI taken up by synchronized iTTFs over 48 h. Meta2d analysis indicates a circadian rhythm of periodicity of 23.8 h. Bars show mean ± s.e.m. of N = 3 per time point. E. Percentage of Cy3-colI taken up by synchronized iTTFs, corrected to the maximum percentage uptake of the time course (pink, bars show mean ± s.e.m. of N=3 per time point), compared to the percentage collagen fibril count over time, corrected to the maximum percentage fibril count of the time course (black, fibrils scored by two independent investigators. Bars show mean± s.e.m. of N=2 with n = 6 repeats at each time point).

### Collagen-I endocytosis is circadian clock regulated, and recycling alone can generate fibrils

Previously, we have shown that clock-synchronized fibroblasts synthesize collagen fibrils in a circadian rhythmic manner [10]; the results here thus far indicate an involvement of endocytosis in collagen fibrillogenesis. Therefore, we hypothesized that collagen-I endocytosis may also be circadian clock regulated. Time-series flow cytometry analyses of fibroblasts incubated with Cy3-colI revealed that the level of Cy3-colI endocytosed by the cells is rhythmic, with a periodicity of 23.8hrs as determined by MetaCycle analysis (**Figure 2D**). When corrected to the running average, the rhythmic nature of Cy3-colI uptake is accentuated (**Figure S2F)**. We noted that, when compared to the number of fibrils produced over time, the peak time of uptake happens before peak fibril numbers (**Figure 2E**). These data suggest that the cells may be endocytosing exogenous collagen under circadian control and holding it in the endosomal compartment, then trafficking the collagen to the plasma membrane for fibril formation.

To eliminate the possibility that the fluorescent fibril-like structures were due to attachment of fluorescently-labeled collagen protomers to pre-existing fibrils already deposited by cells, fibrils already deposited by cells, we performed an siRNA-mediated knockdown against *Col1a1* (siCol1a1) to target endogenous collagen production. Fibroblasts were then analyzed by IF using anti-collagen-I or anti-fibronectin (FN1) antibodies. Control (scrambled siRNAs, scr) cells synthesized collagen-I and fibronectin (**Figure 3A**, top row) with defined fibrillar structures. In contrast, cells treated with siCol1a1 synthesized lower-levels of collagen-I, with significantly few collagen-I fibrillar structures as well as lower intracellular collagen-I signal (**Figure 3A**, bottom row; quantification plots). A ~ 90% reduction in *Col1a1* mRNA was confirmed using qPCR (**Figure S3A**).

**Figure 3:**
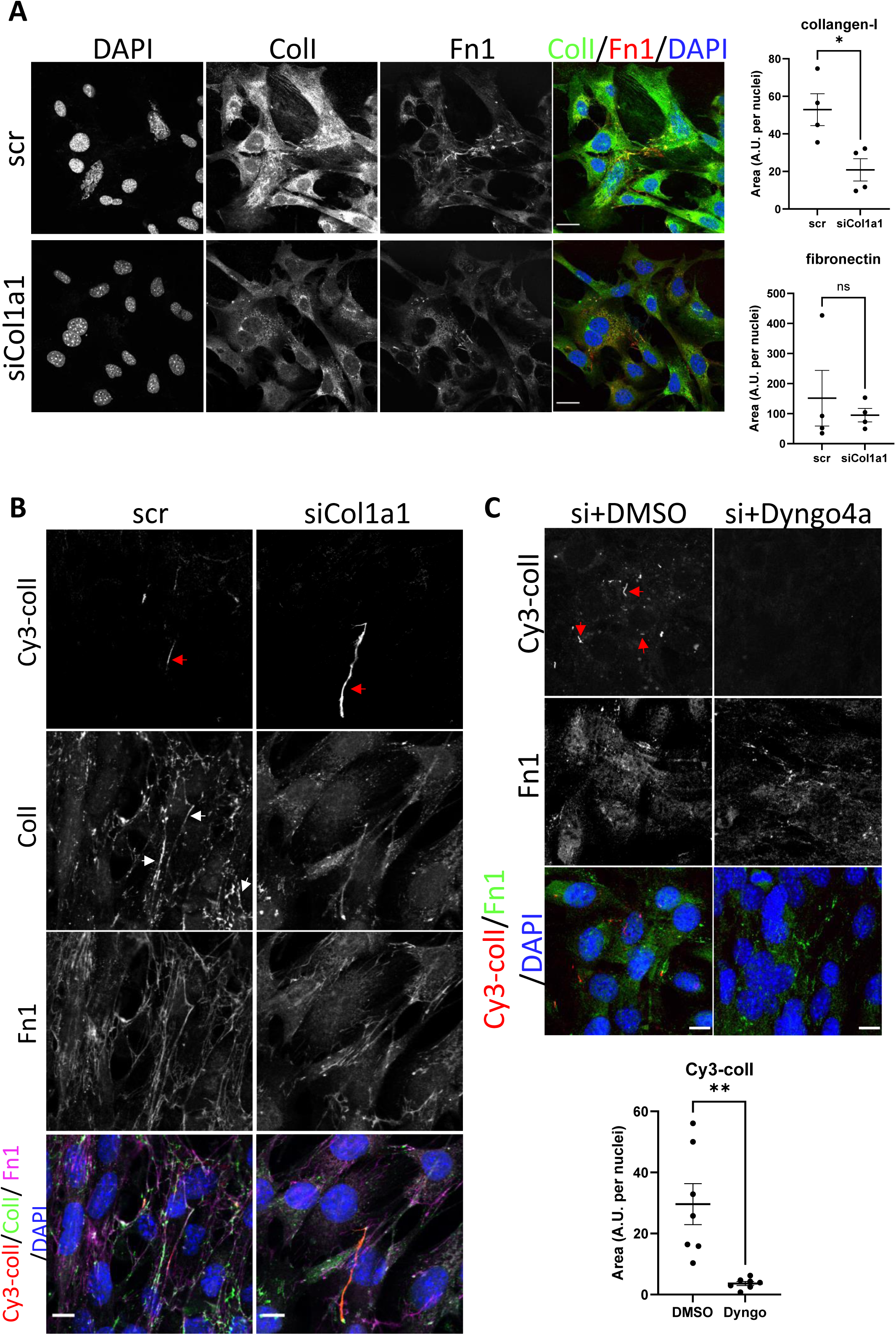
Collagen-I recycling can generate fibrils. A. Fluorescent image series of iTTFs treated with scrambled control (top panel, scr), and siRNA against col1a1 (bottom panel, siCol1a1). Labels on top denotes the fluorescence channel corresponding to proteins detected (ColI – collagen-I Fn1 – fibronectin);. Quantification of collagen-I and fibronectin signal to the right. Representative of N = 4. Scale bar = 25 µm. * *p =* 0.021. B. Fluorescent image series of scr (left column) and siCol1a1 (right column) iTTFs incubated with Cy3-colI. Labels on left denotes the fluorescence channel(s) corresponding to proteins detected (ColI – collagen-I Fn1 – fibronectin). Cy3-colI fibrils highlighted by red arrows, and collagen-I fibrils highlighted by white arrows. Both scr cells and siCol1a1 cells can take up exogenous collagen-I and recycle to form collagen-I fibril. Representative of N>3. Scale bar = 10 *μ*m. C. Fluorescent image series of siCol1a1 iTTFs treated with DMSO control (left) or Dyngo4a (right) during Cy3-colI uptake, followed by further culture for 72 h. Labels on left denotes the fluorescence channel corresponding to proteins detected (ColI – collagen-I Fn1 – fibronectin). Quantification of Cy3-colI signal to the bottom. Dyngo4a treatment led to a reduction of Cy3-colI fibrils. Representative of N>3. Scale bar = 20 µm. ** *p =* 0.0022

Flow cytometry analysis of cells incubated with Cy3-colI revealed that siCol1a1 fibroblasts retained the ability to endocytose exogenous collagen (**Figure S3B**). To assess if siCol1a1 fibroblasts can assemble a collagen fibril with exogenous collagen, Cy3-colI was incubated with scr or siCol1a1 cells for 1 hour, before trypsinization and replating to ensure no cell surface associated Cy3-colI remains. Within 3 days, Cy3-positive collagen fibrils were observed in both scr (**Figure 3B**, left column) and siCol1a1 cells (**Figure 3B**, right column). The observation of fibrillar Cy3 signals in siCol1a1 cells showed that the cells can repurpose collagen into fibrils without the requirement for intrinsic collagen-I production (red arrow, **Figure 3B**). The cells were also probed for collagen-I using an anti-collagen-I antibody with a secondary antibody conjugated to Alexa-488 (**Figure 3B**, second row). In scr cells we detected multiple fibrillar structures that did not contain Cy3-colI (**Figure 3B**, white arrows), indicating fibrils derived from endogenous collagen-I. In the siCol1a1 cells we detected a more diffuse signal and puncta within the cell, but no discernible Alexa-488-only collagen-I fibrils (**Figure 3B**). These results indicate that fibroblasts can assemble collagen-I fibrils from recycled exogenous collagen-I. Importantly, siCol1a1 cells treated with Dyngo4a could not effectively form Cy3-collagen-containing fibrils under these conditions, indicative of a requirement for endocytic-recycling of exogenous collagen-I for fibrillogenesis, when endogenous levels are insufficient (**Figure 3C**). To confirm these findings, we isolated primary tendon fibroblasts from the Col1a2-CreERT2::Col1a1-fl/fl (termed CKO) mouse that had either been treated with tamoxifen (CKO+) to produce fibroblasts that cannot synthesise collagen-I or from untreated mice to yield matched controls (CKO-). Fibroblasts were then analysed by immunofluorescence microscopy using anti-collagen-I antibodies. As expected CKO-cells synthesised collagen-I, and CKO+ cells showed no collagen-I expression (**Figure 4A**). Quantitative PCR analysis of the cells confirmed the absence of Col1a1 mRNAs containing exon 6 to exon 8 sequences, as expected from the location of the LoxP sites in the Col1a1 gene (**Figure S3C**).

**Figure 4:**
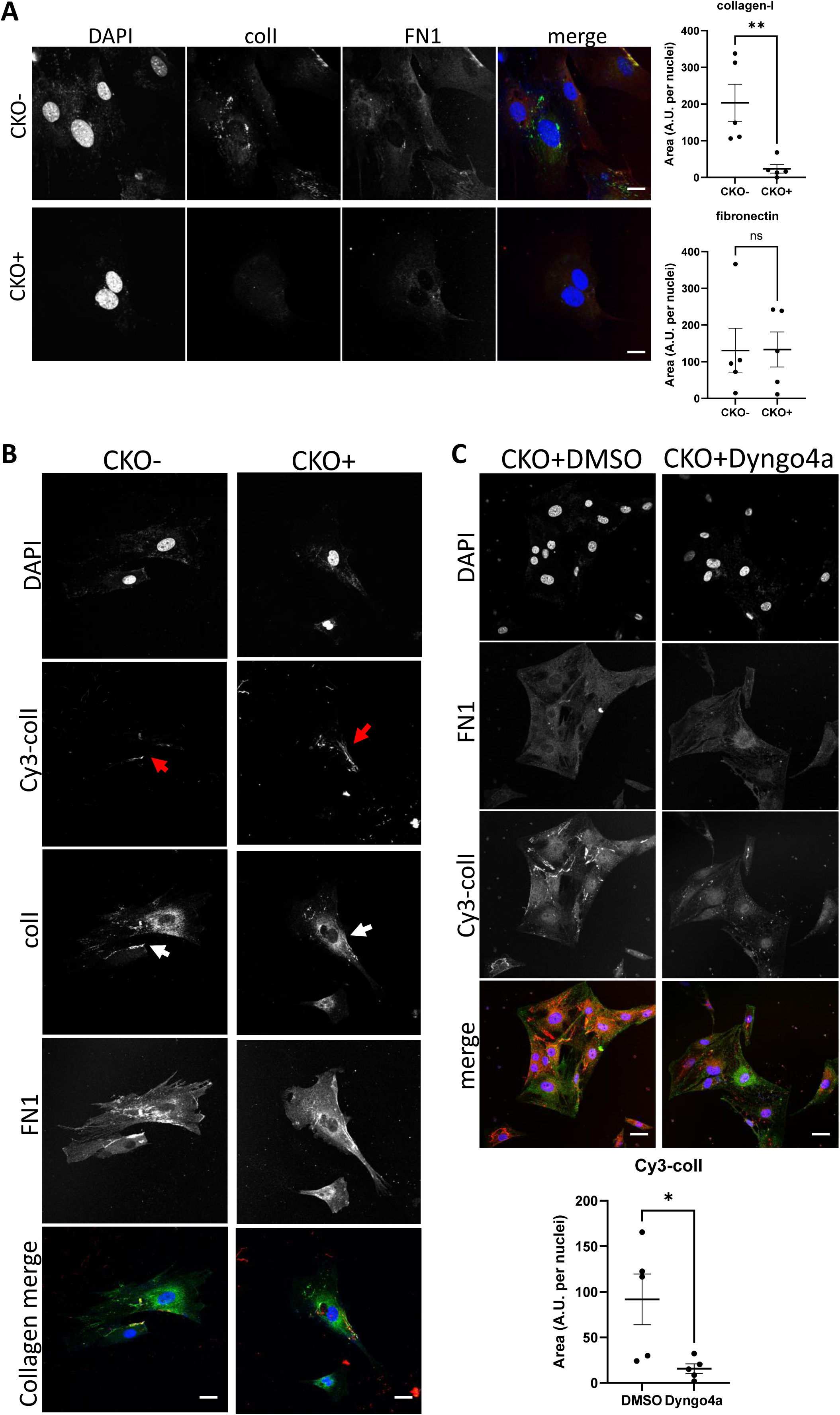
Fibroblasts without endogenous collagen-I can effectively make fibrils by endocytic-recycling of exogenous collagen. A. Fluorescent images of primary tail tendon fibroblasts isolated from control mice (top panel, CKO-), and tamoxifen-treated collagen-knockout mice (bottom panel, CKO+). Labels on top denotes the fluorescence channel corresponding to proteins detected. Quantification of collagen-I and fibronectin fluorescence signal to the right. Representative of N=3. Scale bar = 10*μ*m. ** *p =* 0.0084. B. Fluorescent images of CKO-/CKO+ tail tendon fibroblasts incubated with Cy3-colI. Labels on top denotes the fluorescence channel corresponding to proteins detected. Cy3-colI fibril highlighted by red arrows, and collagen-I fibril highlighted by white arrows. Both CKO- and CKO+ cells can take up exogenous collagen-I and recycle to form collagen-I fibrils. Representative of N>3. Scale bar = 25*μ*m. C. Fluorescent image series of CKO+ tail tendon fibroblasts treated with DMSO control (left) or Dyngo4a (right) during Cy3-colI uptake, followed by further culture for 72 h. Labels on left denotes the fluorescence channel corresponding to proteins detected. Quantification of Cy3-colI signal to the bottom. Dyngo4a treatment led to a significant reduction of Cy3-colI fibrils. Representative of N>3. Scale bar = 25 µm. * *p =* 0.00273.

Flow analysis of cells incubated with Cy3-colI revealed collagen knockout fibroblasts retained the ability to endocytose labelled collagen (**Figure S3D**). To assess whether collagen knockout fibroblasts were able to assemble a collagen fibril, Cy3-colI was incubated with cells before the cells were released by trypsin and replated, to ensure that Cy3 signals arose only from Cy3-colI endocytosed by cells. Within 3 days, Cy3-positive collagen fibres could be observed in both CKO- (**Figure 4B, left column**) and CKO+ cells (**Figure 4B, right column**). The observation of fibrillar Cy3 signals in CKO+ cells showed that the cells can repurpose collagen into fibrils without the requirement for intrinsic collagen-I (**red arrows, Figure 4B**). The cells were also probed for collagen-I using an anti-collagen-I antibody with a secondary antibody conjugated to Alexa-488 (**Figure 4B, second row**). In CKO-cells we detected a diffuse signal over the entire cell and an indication of fibrillar structures. In the CKO+ cells we detected fibrillar structures at the periphery of the cell, and the same diffuse signal seen in CKO-cells (**Figure 4B, white arrows**). We suspect that the diffuse signal observed in CKO+ cells is due to incomplete labelling of collagen-I in our preparation of Cy3-colI, as complete labelling of lysine residues would cause collagen-I to be unable to form fibrils [25]. Nonetheless, Dyngo4a treatment led to a lack of Cy3-colI fibrils (**Figure 4C, right panel and quantification plot)**; thus, these results demonstrated that fibroblasts can effectively form fibrils from exogenous collagen alone.

### VPS33B controls collagen fibril formation but not protomeric secretion

VPS33B is situated in a post-Golgi compartment where it is involved in endosomal trafficking with multiple functions including extracellular vesicle formation [29], modulation of p53 signaling [30], and maintenance of cell polarity [31], dependent on cell type. We previously identified VPS33B to be a circadian-controlled component of collagen-I homeostasis in fibroblasts [10]. As a result, we decided to further investigate its role in endosomal recycling of collagen-I.

We confirmed our previous finding that CRISPR-knockout of VPS33B (VPSko, as verified by western blot analysis and qPCR analysis, **Figure S4A and B**) in tendon fibroblasts led to fewer collagen fibrils without impacting proliferation (**Figure S4C**), as evidenced by both electron microscopy (**Figure 5A**) and IF imaging (**Figure 5B**). Decellularized matrices showed that VPSko fibroblasts produced less matrix by mass than control, which was mirrored by a reduction in hydroxyproline content (**Figure 5C**). We then stably overexpressed VPS33B in fibroblasts (VPSoe), as confirmed by western blot, qPCR analyses, and flow cytometry of transfected cells (**Figure S4D-F**). IF staining indicated a greater number of collagen-I fibrils in VPSoe cells (**Figure 5D**), although the mean total matrix and mean total hydroxyproline in VPSoe cultures was not significantly higher than control (**Figure 5E**). As VPSoe cells showed equivalent proliferation to controls (**Figure S4G**), this suggests VPSoe specifically enhanced the assembly of collagen fibrils. We then performed time-series IF on synchronized cell cultures to quantify the number of collagen fibrils formed. Tendon fibroblasts exhibited a ~ 24 h rhythmic fluctuation in collagen-I fibril numbers, whereas VPSoe cells continuously deposited collagen-I fibrils over the 55-h period (**Figure 5F**). MetaCycle analyses indicated that fluctuation of fibril numbers in control and VPSoe cultures occurred at a periodicity of 22.7 h and 28.0 h respectively (**Figure S4H**), indicating that continuous expression of VPS33B leads to loss of collagen-I fibril circadian homeostasis. Interestingly, when assessing the levels of secreted soluble collagen-I protomers from control and VPSoe cells, VPSoe CM have lower levels of soluble collagen-I (**Figure 5G**). In addition, when VPS33B is knocked-down using siRNA, there is an elevation of soluble collagen-I secreted; siVPS33B on VPSoe cells also increased the levels of collagen-I in CM (**Figure S4I, J**), whilst VPS33B levels are not correlated with *Col1a1* expression levels (**Figure S4K**). This finding indicates that VPS33B is specifically involved in collagen-I fibril assembly, and not secretion or translation. The reduction of secreted soluble collagen-I in VPSoe cells further supports that VPS33B directs collagen-I towards fibril assembly.

**Figure 5:**
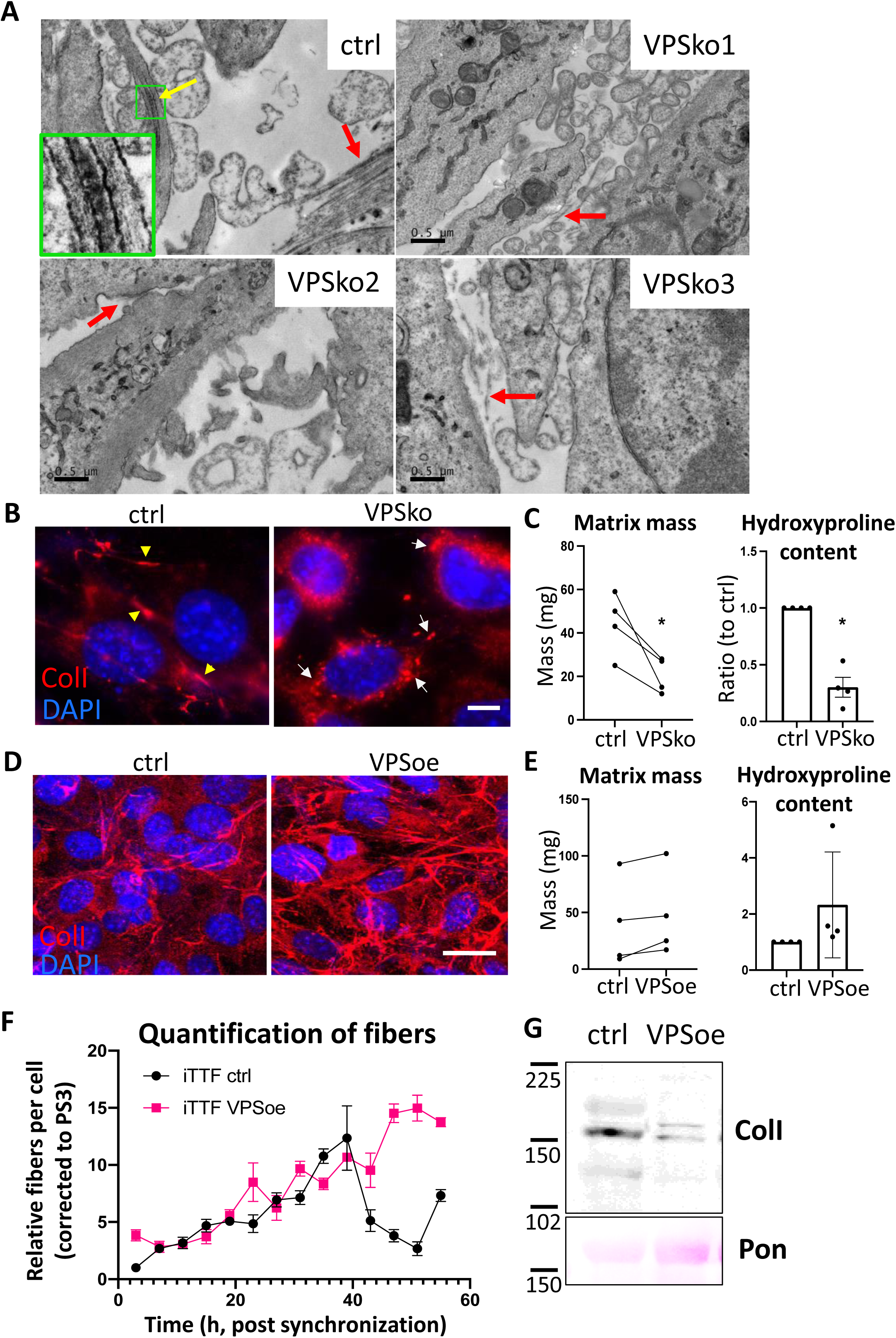
VPS33B controls collagen fibril formation at the plasma membrane in a rhythmic manner. A. Electron microcopy images of fibroblasts plated on Aclar and grown for a week before fixation and imaging. Ctrl culture has numerous collagen-I fibrils, as pointed out by arrows. Yellow arrow points to a fibripositor, and green box is expanded to the left bottom corner, showing the distinct D-banding pattern of collagen-I fibril when observed with electron microscopy. VPSko clones all have fewer and thinner fibrils present in the culture (pointed out by red arrows). Representative of N = 3. Scale bar = 0.5 µm. B. Fluorescence images of collagen-I (red) and DAPI counterstain in ctrl and VPSko iTTFs. Yellow arrows indicating collagen fibrils, and white arrows pointing to collagen-I presence in intracellular vesicles. Representative of N>6. Scale bar = 25 µm. C. Matrix deposition by ctrl or VPSko iTTFs, after one week of culture. Left: decellularized matrix mass. N = 4, * *p =* 0.0299. Right: hydroxyproline content presented as a ratio between ctrl and VPSko cells. N = 4, \**p =* 0.0254. Ratio-paired t-test used. D. Fluorescence images of collagen-I (red) and DAPI counterstain in ctrl and VPSoe iTTFs. Representative of N>6. Scale bar = 20 µm. E. Matrix deposition by ctrl or VPSoe iTTFs, after one week of culture. Left: decellularized matrix mass, N = 4. Right: hydroxyproline content presented as a ratio between ctrl and VPSoe cells, N = 4. Ratio-paired t-test used. F. Relative collagen fibril count in synchronized ctrl (black) and VPSoe (pink) iTTFs, corrected to the number of fibrils in ctrl cultures at start of time course. Fibrils scored by two independent investigators. Bars show mean± s.e.m. of N = 2 with n=6 at each time point. G. Western blot analysis of conditioned media taken from ctrl and VPSoe iTTFs after 72 h in culture. Top: probed with collage-I antibody (ColI), bottom: counterstained with Ponceau (Pon) as control. Protein molecular weight ladders to the left (in kDa). Representative of N = 3.

### VPS33B-positive intracellular structures contain collagen-I

We then utilized the split-GFP system [32, 33] to investigate how VPS33B specifically directs collagen-I to fibril assembly. Here, a GFP signal will be present if VPS33B co-traffics with collagen-I. We, and others, have previously demonstrated that insertion of tags at the N-terminus of proα2(I) or proα1(I) chain does not interfere with collagen-I folding or secretion [34, 35]. To determine which terminus of the VPS33B protein GFP1-10 should be added, we performed computational ΔG analysis [36], which predicts that VPS33B contains two regions of extended hydrophobicity towards its C-terminus (**Figure S5A**). The first region (residues 565-587, denoted TMD1) has a ΔG of −0.62 kcal/mol, consistent with a single pass type IV transmembrane domain (TMD), known as a tail-anchor. This arrangement locates the short C-terminus of VPS33B inside the lumen of the endoplasmic reticulum (ER), and subsequently within the endosomal compartment [37]. The second region (residues 591-609, denoted HR1) is significantly less hydrophobic, with a ΔG of +2.61 kcal/mol (**Figure S5B**). Its presence raises the possibility that VPS33B may contain two TMDs that, depending on their relative membrane topologies, result in either a luminal or cytosolic C-terminus (**Figure S5C**).

To define the topology of VPS33B, we used a well-established *in vitro* system where newly synthesized and radiolabeled proteins of interest are inserted into the membrane of ER-derived canine pancreatic microsomes, and created constructs with ER luminal modification of either endogenous N-glycosylation sites (N54, N545 in VPS33B), or artificial sites in an appended OPG2 tag (N2, N15 in residues 1-18 of bovine rhodopsin, Uniprot: P02699) as a robust reporter for membrane protein topology in the ER (**Figure S5D, E**) [38]. Due to the N-terminal region of VPS33B being highly aggregation-prone in our *in vitro* system (**Figure S5F, G**), we created three additional chimeric proteins comprised of different regions of VPS33B and Sec61β to investigate the topology of VPS33B (**Figure S5H**). Whilst a small proportion of each chimera continued to pellet when synthesized in the absence of ER-derived microsomes (**Figure S5I,** lanes 3, 6, 9), N-glycosylated species were now clearly identifiable for each of the chimeras tested.

In chimera 1, given the respective efficiency of membrane insertion of the TA region and N-glycosylation of the C-terminal OPG tag of Sec61βOPG2 (**Figure S5I,** lanes 1-2), we attribute the N-glycosylated species to the C-terminal translocation of OPG2 tag, although it is evident that the remaining short stretch of VPS33B (residues 414-564) still impedes efficient membrane insertion, likely due to aggregation (**Figure S5I**, lanes 3-5) [39, 40]. For chimera 2, the efficient modification of its distal N-glycosylation site (N15 of the OPG2 tag) but inefficient use of the proximal site (N2 of the OPG2 tag) reflects the latter residues’ close proximity to the ER membrane [41], and supports the *bona fide* membrane insertion of the VPS33B TMD1 and thus translocation of the C-terminal OPG2 tag into the ER lumen (**Figure S5I**, lanes 6-8). Interestingly, the inclusion of HR1 to the C-terminus of TMD1 in chimera 3 results in a substantial qualitative reduction in the amount of protein that is N-glycosylated (**Figure S5I**, lanes 7 and 10). Further, for the fraction of chimera 3 that is modified, the majority is doubly N-glycosylated; most likely due to the extra length provided by HR1, resulting in both N-glycosylation sites on the OPG2 tag (N2 and N15) now being efficiently modified [41].

The clear reduction in the proportion of N-glycosylated species obtained with chimera 3 compared to chimera 2 (**Figure S5I**) indicates that only a small proportion of HR1 and the appended OPG2 tag are successfully translocated into the ER lumen. This is likely due to a combination of the low proportion of hydrophobic residues and the presence of several charged and polar amino acids in HR1[42]. We thus propose that, for a minority of VPS33B, TMD1 is inserted into the ER membrane as a tail-anchored region with a luminal HR1 that most likely remains associated with the inner leaflet of the bilayer (N-cytosolic, C-luminal; **Figure 6A**, right). In contrast, the majority of VPS33B likely assumes a ‘hairpin’ like conformation in the ER membrane where both its N- and C-termini remain in the cytosol (N-cytosolic, C-cytosolic) with either: i) a partially membrane inserted TMD1, with HR1 associated with the outer leaflet of the ER membrane (**Figure 6A**, left); or ii) a fully membrane inserted TMD1 followed by the marginally hydrophobic HR1 which may span the membrane through the formation of stabilizing hydrogen bonds with TMD1 (**Figure 6A**, middle) [43].

**Figure 6:**
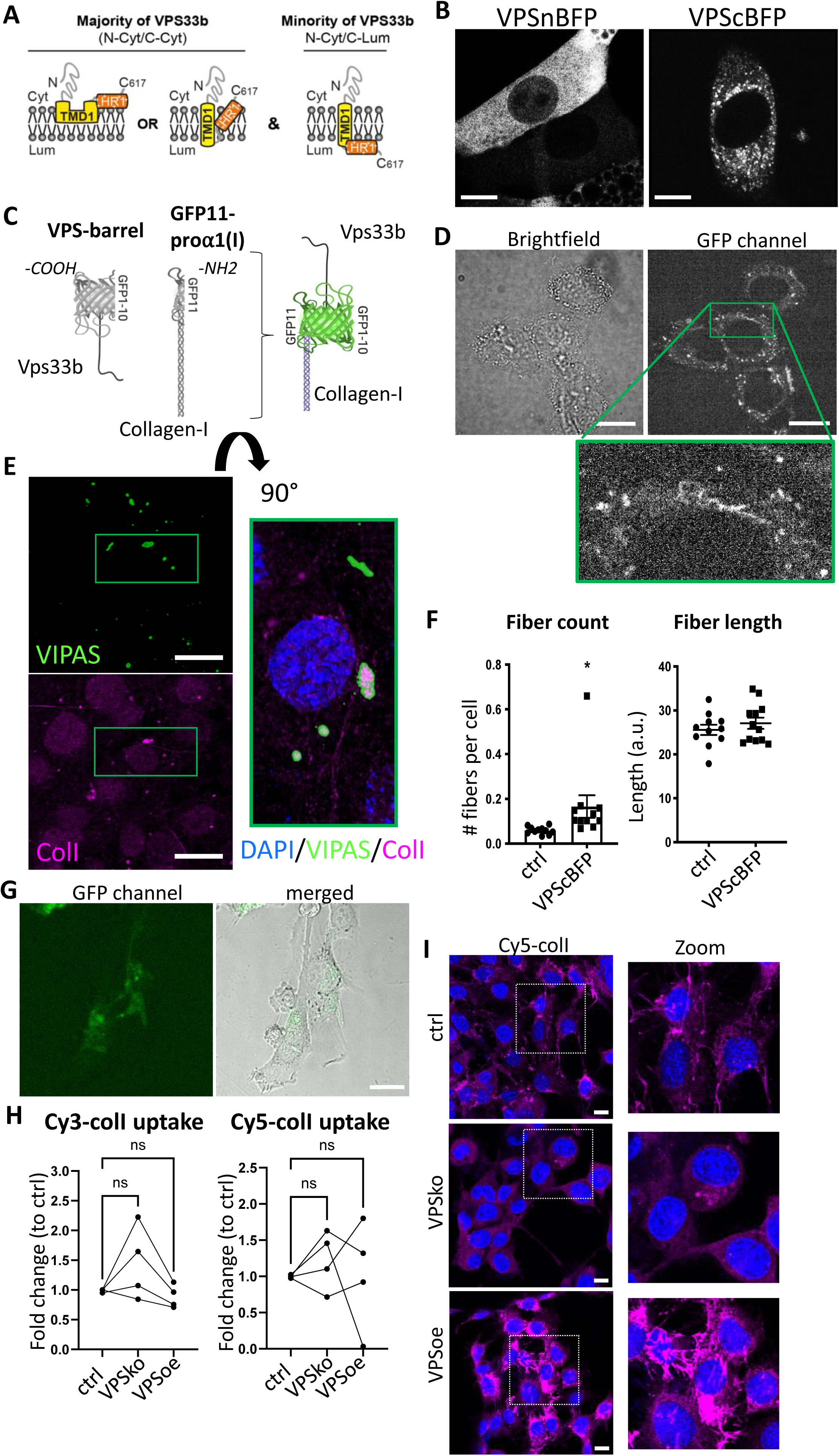
Procollagen-I and VPS33B localize to the same compartments. A. Schematic depicting the proposed membrane topologies of VPS33b. B. iTTFs expressing BFP-tagged VPS33B. Left: BFP tagged on the N-terminal end of VPS33B (VPSnBFP), Right: BFP tagged on the C-terminal end (VPScBFP). Images taken in Airy mode. Representative of N>4. Scale bar = 10 µm. C. Schematic of the split-GFP system. GFP1-10 barrel is introduced into VPS33B (VPS-barrel), and GFP11 to alpha-1 chain of Collagen-I (GFP11-pro*α*1(I)). If the two tagged proteins co-localize (e.g. in a vesicle), a GFP signal will be emitted. D. Brightfield (top) and fluorescence (middle) images of iTTFs expressing both VPS-barrel and GFP11-pro*α*1(I) constructs. Representative of N=5. Green box is expanded to the bottom, to highlight the punctate fluorescence signals within intracellular vesicular structures, as well as fibril-like structures suggestive of fibril assembly sites. Scale bar = 20 µm. E. Fluorescence images of VIPAS (green), collagen-I (red) and DAPI counterstain in iTTFs. Representative of N=3. Green box is expanded to the right (flipped 90°) to show strong VIPAS signal encasing collagen-I. Scale bar = 25 µm. F. Quantification of average number of fibrils per cell (left) and average fibril length (right) in control endogenously-tagged Dendra-colI expressing 3T3 cells (ctrl) and Dendra-colI expressing 3T3 overexpressing VPScBFP (VPScBFP). >500 cells quantified per condition. N=12. \**P*=0.048. G. Brightfield (left) and fluorescence (middle) images of iTTFs expressing VPS-barrel incubated with conditioned media containing Col1a1-GFP11 for 24hr. Scale bar = 25*μ*m. H. Line charts comparing the percentage of iTTFs that have taken up 5 µg/mL Cy3-colI (left) and Cy5-colI (right) after one hour incubation between control (ctrl), VPS33B-knockout (VPSko) and VPS33B-overexpressing (VPSoe) cells, corrected to control. RM one-way ANOVA analysis was performed. N = 4. I. Fluorescence images of iTTFs of different levels of VPS33B expression, fed with Cy5-colI and further cultured for 72 h. Cultures were counterstained with DAPI. Box expanded to right of images to show zoomed-in images of the fibrils produced by the fibroblasts. Representative of N = 2.

To test our topology findings, we inserted BFP at either the N- (VPSnBFP) or C-terminus (VPScBFP) of VPS33B. In cells expressing VPSnBFP, the fluorescence signal was diffuse throughout the cell body, and in some cases appeared to be completely excluded from circular structures (**Figure 6B**, left), suggestive of protein mistargeting. In contrast, VPScBFP-expressing cells have punctate blue fluorescence signals and peripherally blue structures (**Figure 6B**, right), supportive of the topology findings. Previous studies have also tagged VPS33B at the C-terminus [11]. Thus, the 214-residue N-terminal fragment (GFP1-10) was cloned onto VPS33B, and the 17-residue C-terminal peptide (GFP11) was cloned onto the proα1(I) chain of collagen-I as previously described [34] (GFP11-proα1(I), **Figure 6C**).

Fibroblasts stably expressing VPS-barrel and GFP11-proα1(I) were imaged to identify any GFP fluorescence. The results showed puncta throughout the cell body (**Figure 6D**). Intriguingly, GFP fluorescence was also observed at the cell periphery (**Figure 6D**, green box zoom). IF staining of endogenous VIPAS39 (a known VPS33B-interacting partner [11]) also revealed co-localization of VIPAS39 with collagen-I in intracellular punctate structures, where in some of the co-localized puncta, the signal of VIPAS39 is strongest surrounding collagen-I (**Figure 6E**, zoom), suggesting that VIPAS39 is encasing collagen-I, not within the lumen but present in proximity with the external membrane of these structures. VPS33B has been demonstrated to interact with VIPAS in regions before the TMD1 site [44]; thus, in all suggested VPS33B topologies herein, VIPAS39 will still be able to interact with VPS33B, and encase collagen-I-containing intracellular structures.

The relationship between VPS33B levels and collagen fibril numbers was confirmed using endogenously-tagged Dendra2-collagen-I expressing 3T3 cells [45], where notably the number of Dendra2-positive fibrils significantly increased when VPS33BcBFP was expressed; however, the average length of the fibril was not significantly different, suggesting that VPS33B is important in fibril initiation but not elongation (**Figure 6F**). Incubation of iTTFs expressing only the VPS-barrel with CM collected from iTTFs expressing only GFP11-proα1(I) revealed that the GFP signal was detected only in intracellular structures and not along the periphery of the cells, where endocytosis takes place (**Figure 6G**). Flow cytometry analyses of fluorescently labelled-colI endocytosed by control, VPSko, and VPSoe fibroblasts also revealed no consistent change in uptake by VPSko or VPSoe cells (**Figure 6H**). Taken together, these results suggest that VPS33B interacts with endocytosed collagen-I within the cell and and trafficks with collagen-I to the extracellular space, and is not involved with collagen-I endocytosis. VPSko cells replated after Cy5-colI uptake have conspicuously fewer fibrils when compared to control. In contrast, VPSoe cells have shorter but more Cy5-colI fibrils (**Figure 5I**), highlighting the role of VPS33B in recycling endocytosed collagen-I to initiate collagen fibrillogenesis.

### Integrin chain α11 mediates VPS33B-dependent fibrillogenesis

Having identified VPS33B as a driver for collagen-I fibril formation but not protomeric secretion, we used biotin cell surface labeling coupled with mass spectrometry protein identification to identify other proteins that may be involved in this process at the cell surface. VPSko and VPSoe fibroblasts were analyzed using a “shotgun” approach. In total, 4121 proteins were identified in total lysates (**Table S1**), and 1691 proteins in the enriched-for-surface-protein samples (**Table S2**). Gene Ontology (GO) Functional Annotation analysis [46, 47] identified the top 25 enriched terms (based on p-values) with the top 5 terms all associated with “extracellular” or “cell surface” (**Figure 7A**), indicative that the biotin-labeling procedure had successfully enriched proteins at the cell surface interacting or associated with the extracellular matrix (ECM).

**Figure 7:**
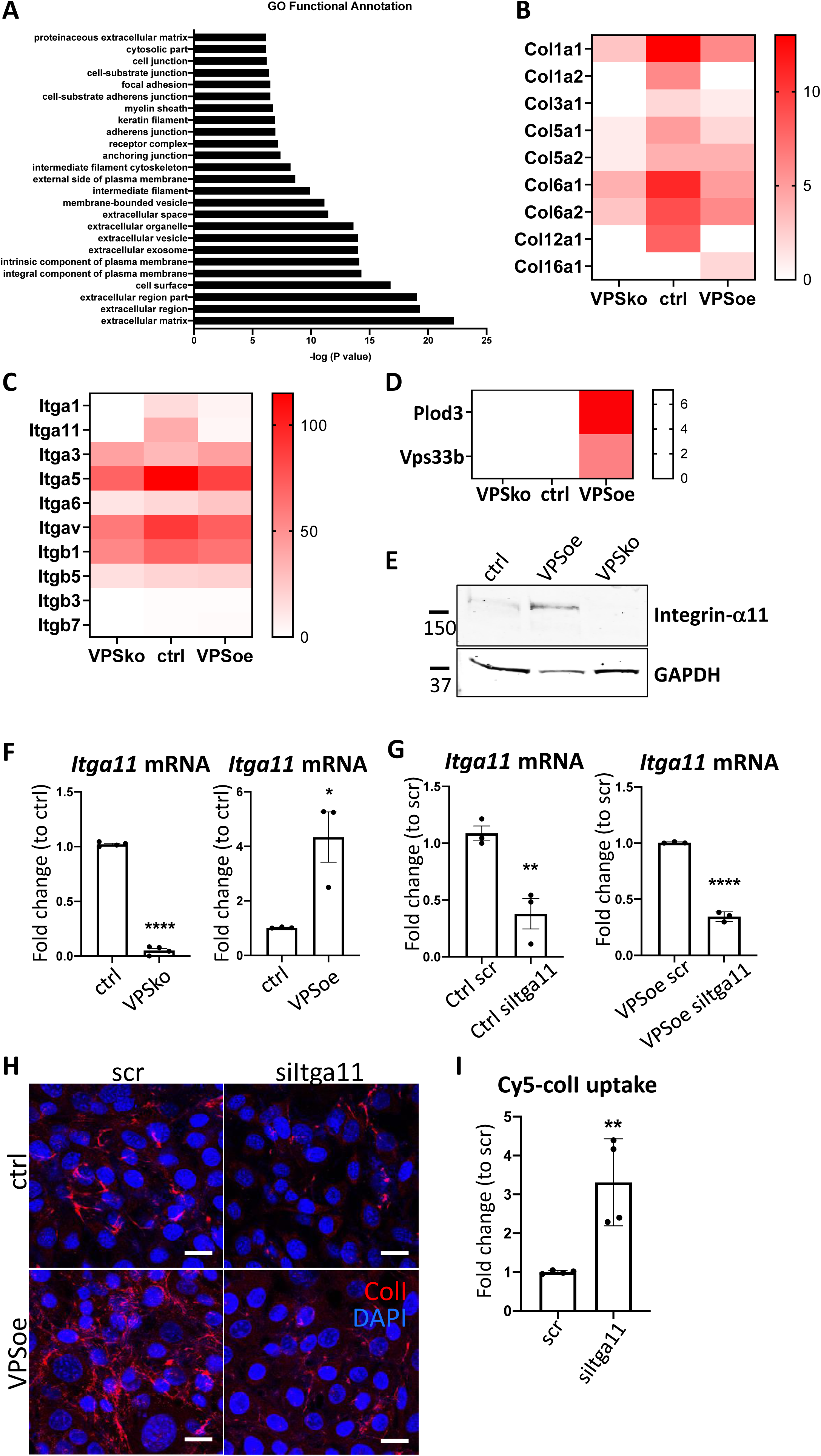
Integrin α11 subunit mediates VPS33B-effects and is required for collagen-I fibrillogenesis. A. Top 25 Functional Annotation of proteins detected in biotin-enriched samples when compared to non-enriched samples based on p-values. Y-axis denotes the GO term, X-axis denotes –log (P value). B. Heatmap representation of spectral counting of collagens detected in biotin-enriched surface proteins from control (ctrl), VPS33B-knockout (VPSko), and VPS33B-overexpressing (VPSoe) iTTFs. Scale denotes quantitative value as normalized to total spectra, as determined by Proteome Discoverer. C. Heatmap representation of spectral counting of integrins detected in biotin-enriched surface proteins from control (ctrl), VPS33B-knockout (VPSko), and VPS33B-overexpressing (VPSoe) iTTFs. Scale denotes quantitative value as normalized to total spectra, as determined by Proteome Discoverer. D. Heatmap representation of spectral counting of Plod3 and VPS33B detected in biotin-enriched surface proteins from control (ctrl), VPS33B-knockout (VPSko), and VPS33B-overexpressing (VPSoe) iTTFs. Scale denotes quantitative value as normalized to total spectra, as determined by Proteome Discoverer. E. Western blot analysis of integrin α11 subunit levels in control (ctrl), VPS33B-overexpressing (VPSoe), VPS33B-knockout (VPSko) iTTFs. Top: probed with integrin α11 antibody, bottom: reprobed with GAPDH antibody. Protein molecular weight ladders to the left (in kDa). Representative of N = 3. F. qPCR analysis of *Itga11* transcript levels in ctrl compared to VPSko iTTFs (left), and ctrl compared to VPSoe iTTFs (right). N>3, \*\*\*\**P<*0.0001, \**P* = 0.0226. G. qPCR analysis of *Itga11* mRNA expression in ctrl (left) or VPSoe (right) iTTFs treated either with scrambled control (scr) or siRNA against Itga11 (siItga11), collected after 96 h. N=3, \*\**P*=0.0091, \*\*\*\**P*<0.0001. H. IF images of ctrl and VPSoe iTTFs treated either with control siRNA (scr) or siRNA again Itga11 (siItga11), after 72 h incubation; collagen-I (red) and DAPI (blue) counterstained. Representative of N = 3. Scale bar = 25 µm. I. Bar chart comparing the percentage of iTTFs that have taken up 5 µg/mL Cy5-colI after one hour incubation between fibroblasts treated with scrambled control (ctrl) or siRNA against Itga11 (siItga11), corrected to scr. N=3. \*\**P*=0.0062.

We then interrogated the differences between VPSko and control cells, visualizing the results in a semi-quantitative manner using spectral counting (**Table S3**). Conspicuously, collagen α1(I) and α2(I) chains were detected at a reduced level at the surface of VPSko cells (**Figure 7B**). The reduction of α1(V) and α2(V) chains (which constitute type V collagen) from the cell surface supports the long-standing view that collagen-V nucleates collagen-I containing fibrils. Gene ontology pointed to integral components of the plasma membrane, which included several integrins. Whilst many integrins were detected in all samples there was an absence of integrin α1 and integrin α11 subunit in VPSko cultures (**Figure 7C**). It is well established that integrins are cell-surface molecules that interact extensively with the ECM [48], with integrin α11β1 functioning as a major collagen binding integrin on fibroblasts [23], which is expressed during development, and is upregulated in subsets of myofibroblasts in tissue and tumor fibrosis [5, 49, 50]. Importantly, whilst the level of integrin β1 chain detected was also reduced in VPSko cultures, it was not as drastic as the reduction of integrin α11 chain. This is likely due to the promiscuous nature of integrin β1 subunit, that is being able to partner with other α subunits for functions other than collagen-I interaction. The absence of integrin α11 subunit from VPSko cells suggested a link between VPS33B-mediated collagen fibrillogenesis and integrin α11β1.

VPS33B was conspicuous by its absence from cell-surface labeling studies of control samples (**Figure 7D**). We have previously struggled to detect VPS33B protein using mass spectrometry [10] and postulate that its absence could be due to low abundance. Regardless, VPS33B was detected at the cell surface in VPSoe samples along with PLOD3 (**Figure 7D**), a lysyl hydroxylase involved in stabilizing collagen and previously identified to be delivered by VPS33B [14]. Complementary western blot analysis of total cell lysates showed that expression of integrin α11 subunit is significantly reduced in VPSko cells and is elevated in VPSoe cells (**Figure 7E**). This correlation between VPS33B and integrin α11 expression levels was confirmed at the mRNA level (**Figure 7F**), inferring a link between VPS33B, collagen fibril, and integrin α11 abundance. We verified this by siRNA knockdown of *Itga11* in control and VPSoe fibroblasts, where knockdown efficiency is confirmed by qPCR (**Figure 7G**). siItga11 in both control and VPSoe cells reduced the number of collagen-I fibrils in culture (**Figure 7H**). Thus, even with elevated levels of VPS33B, integrin α11 subunit is required for collagen-I fibrillogenesis. Interestingly, knocking-down integrin α11 subunit increased exogenous collagen-I uptake as demonstrated by flow cytometry (**Figure 7I**). Thus, we propose that both VPS33B and integrin α11 are involved in directing collagen-I protomers to the formation of collagen-I fibrils, and not mediators of collagen-I endocytosis.

### Integrin α11 subunit is localized to the fibroblastic focus of idiopathic pulmonary fibrosis

We next investigated if VPS33B and integrin α11 are involved in human fibrotic diseases, where accumulation of collagen fibrils is a disease hallmark. Lung fibroblasts isolated from control individuals or individuals suffering from IPF were cultured and the mRNA levels of *COL1A1*, *ITGA11,* and *VPS33B* determined (**Figure 8A**). Although *COL1A1* was similarly expressed, the levels of *VPS33B* and *ITGA11* were significantly increased in IPF fibroblasts compared to control. We next assessed the endocytic capacities of collagen-I in human lung fibroblasts, and confirmed co-localisation of exogenous Cy5-colI with Rab5-GFP (**Figure S6A**). We also found that IPF fibroblasts endocytosed significantly more Cy5-labelled exogenous collagen-I when compared to control fibroblasts (**Figure 8B**, left); uptake of Cy3-labelled exogenous collagen-I was also elevated, albeit not significantly (**Figure 8B**, right). Subsequent culture of cells that have taken up exogenous collagen-I revealed that IPF fibroblasts made significantly more fluorescently-labeled collagen fibrils, indicating enhanced recycling of endocytosed collagen-I to generate new fibrils (**Figure 8C**). The relationship between VPS33B, ITGA11 and endocytic recycling of collagen-I for fibrillogenesis was further confirmed using siRNA (**Figure S6B**), where knock-down of either proteins led to a significant decrease in recycling of exogenous Cy5-ColI (**Figure 8D**). This *in vitro* observation was also represented in IPF pathology, where patient-derived lung samples showed enrichment of collagen-I which overlaps with both integrin α11 subunit and VPS33B within the IPF hallmark lesion (termed the fibroblastic focus [24], encircled by red dotted line, **Figure 9A**). In sites of emerging fibrotic remodeling, integrin α11 subunit and VPS33B are also detected (red asterisks, **Figure 9B**; additional N = 3 IPF specimens, **Figure S7A**), indicating a role for VPS33B/integrin α11 chain in collagen-I deposition in the context of IPF. In control lung samples, collagen-I and VPS33B were present whereas integrin α11 subunit was detected at negligible levels (**Figure 8C, Figure S7B**). These results indicate that the proteins required for assembly of collagen into fibrils (i.e. integrin α11, VPS33B) are present at the fibrotic fronts of IPF, and that enhanced endocytic recycling of collagen-I by fibroblasts may be a disease-potentiating mechanism in IPF.

**Figure 8:**
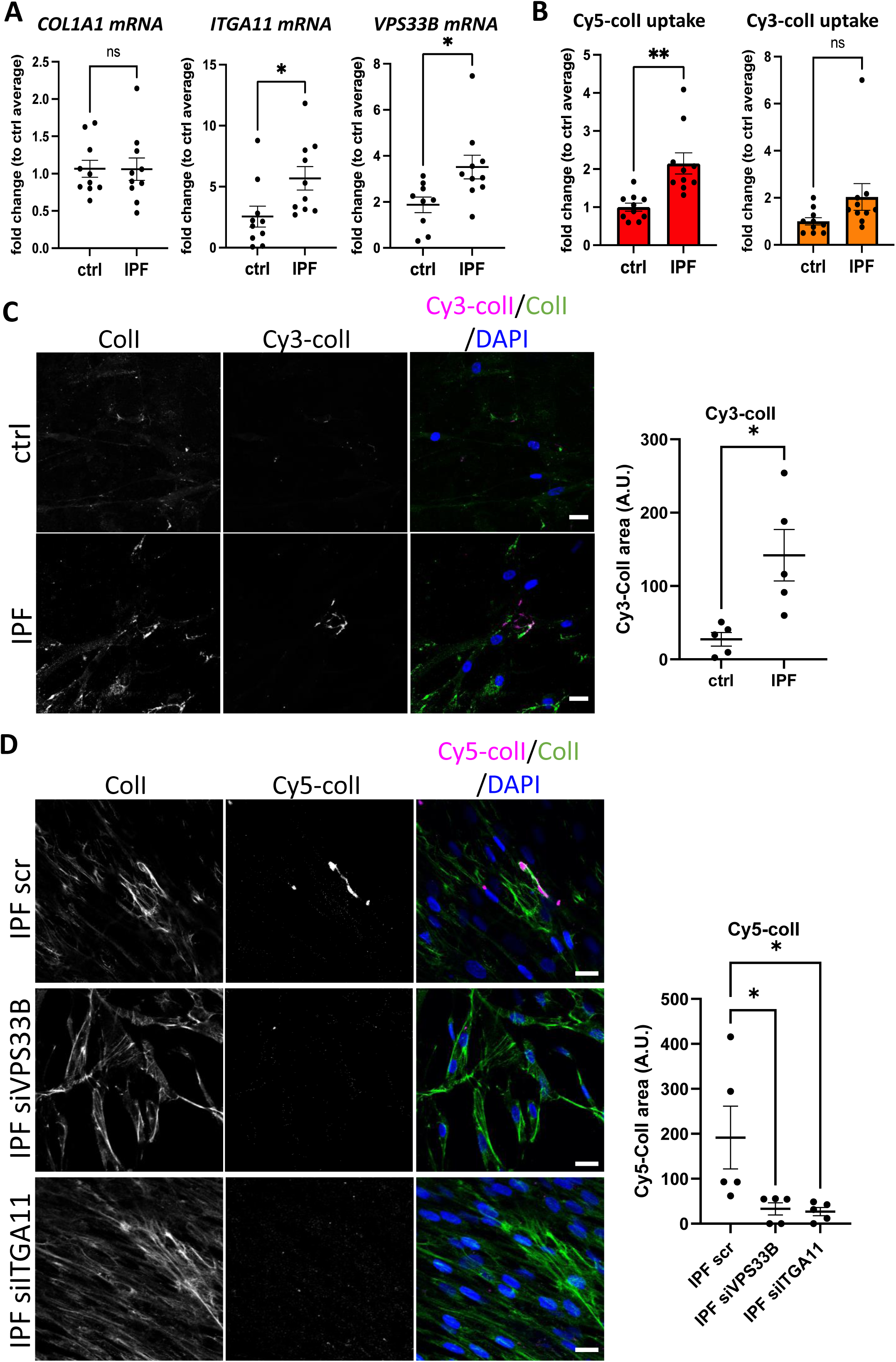
Fibroblasts derived from IPF patients have higher collagen endocytic-recycling capacity that is mediated by VPS33B and ITGA11. A. qPCR analysis of patient-derived fibroblasts isolated from control (ctrl) or IPF lungs. Bars showing mean± s.e.m, 5 patients in each group from 2 independent experiments (technical repeats not shown here). *Itga11, *p* = 0.0259; *VPS33B*, \**p* = 0.0183. B. Fold change of percentage Cy5-colI (left) or Cy3-colI (right) taken up by ctrl or IPF lung fibroblasts, corrected to average of control fibroblasts. Bars showing mean± s.e.m., 5 patients in each group from 2 independent experiments (technical repeats not shown here). \*\**p* = 0.003. C. Fluorescent images of ctrl or IPF lung fibroblasts that have taken up Cy3-colI (magenta), followed by further culture for 48 h in presence of ascorbic acid, before subjected to collagen-I staining (green). Labels on top denotes the fluorescence channel corresponding to proteins detected. Quantification of Cy3-colI signal to the right. IPF fibroblasts produced more Cy3-labelled fibrillar structures. Representative of N = 5. Scale bar = 20 µm. * *p =* 0.0135. D. Fluorescent images of IPF lung fibroblasts treated with siRNA scrambled control (scr), siRNA against VPS33B (siVPS33B) or siRNA against ITGA11 (siITGA11) prior to uptake of Cy5-colI (magenta). This was followed by further culture for 48 h in presence of ascorbic acid, before subjected to collagen-I staining (green). Labels on top denotes the fluorescence channel corresponding to proteins detected. Quantification of Cy5-colI signal to the right. Both siVPS33B and siITGA11 significantly reduced recycled collagen signals. Representative of n = 5 across N = 2. Ordinary one-way ANOVA with multiple comparisions (to scr) was performed on quantification of Cy5-colI signal. siVPS33B, * *p =* 0.0341; siITGA11, * *p =* 0.0282.

**Figure 9:**
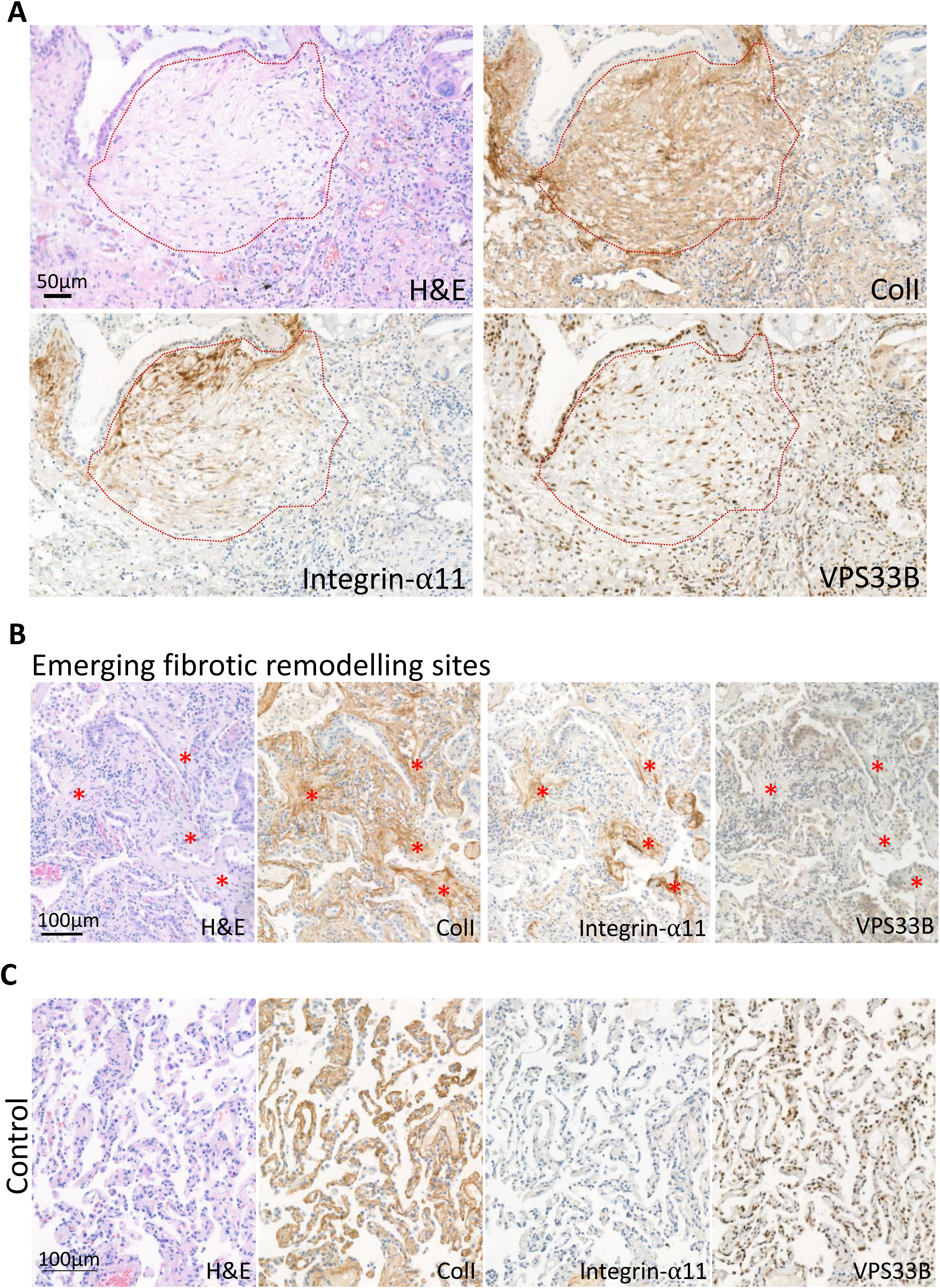
The IPF fibrotic focus is positive for integrin α11 subunit and VPS33B. A. Immunohistochemistry of IPF patient (patient 1) with red dotted line outlining the fibroblastic focus, the hallmark lesion of IPF. Sections were stained with hematoxylin and eosin (H&E), collagen-I (ColI), integrin-α11, VPS33B. Scale bar = 50 µm. B. Immunohistochemistry of IPF patient 4 showing regions of emerging fibrotic remodeling with evidence of fibroblastic foci formation (red asterisks). Sections were stained with H&E, ColI, integrin-α11, VPS33B. Scale bar = 100 µm. C. Immunohistochemistry of 5 µm thick sequential lung sections taken from lungs classified as control (Control 1). Sections were stained with hematoxylin and eosin (H&E), collagen-I (ColI), integrin-α11, VPS33B. Scale bar = 100 µm.

### VPS33B and integrin α11 subunit levels are elevat3ed in chronic skin wounds

Fibrosis has been described as a dysregulation of the normal wound-healing process [51]. Thus, we postulated that chronic skin wounds, a similar chronic inflammation unresolved wound-healing condition, may also share a similar pathological molecular signature as IPF. Immunohistochemical (IHC) staining revealed that in human chronic skin wounds, the expressions of integrin α11 subunit and VPS33B are elevated when compared to normal skin areas taken from the same patient (**Figure 9**), where expression was evident at both the fibrotic wound margins and perivascular tissues, indicative of areas under constant collagen remodeling, suggestive of a similar dysregulation in collagen fibrillogenesis pathway utilizing VPS33B and integrin α11.

## Discussion

In this study we have identified an endocytic recycling mechanism for type I collagen fibrillogenesis that is under circadian regulation. Integral to this process is VPS33B (a circadian clock-regulated endosomal tethering molecule) and integrin α11 subunit (a collagen-binding transmembrane receptor when partnered with integrin β1 subunit). Collagen-I co-traffics with VPS33B to the plasma membrane for fibrillogenesis, which requires integrin α11. These proteins are all enhanced at active sites of collagen pathologies such as fibrosis and chronic skin wounds, suggestive of a common disease mechanism.

Previous research [16, 52–55] has focused on degradation or signaling as the endpoint of collagen-I endocytosis. However, our results show that collagen-I endocytosis contributes towards fibril formation, which is reduced when endocytosis is inhibited. The decision between degrading or recycling endocytosed collagen may depend on cell type, microenvironment, or collagen type itself. In complex environments, for example a wound, degradation of damaged collagen molecules and recycling of structurally sound collagen molecules could be beneficial. A caveat in this present study is the use of Dyngo4a to inhibit endocytosis. It was demonstrated that there may be non-dynamin targeting effects with Dyngo4a when comparing the effects of dynamin triple knock-out and drug treatment (e.g. fluid-phase endocytosis and membrane ruffling) [56]; here we have shown that collagen uptake is likely through receptor-mediated and not fluid-phase endocytosis, although whether periphery membrane ruffling alters collagen deposition remains to be investigated. As total secretion did not appear to be affected by Dyngo4a treatment, as indicated by Ponceau stain on western blots of CM, we take this to suggest that Dyngo4a in this study is not affecting overall secretion and is mostly acting on endocytosis. Further studies targeting multiple endocytosis routes, either through genetic manipulation or drug treatments, will help shed light on how collagen uptake is controlled, and if different forms of collagen (i.e. protomers, fibril) require different mechanisms. Nonetheless, the requirement of VPS33B for fibril assembly confirms the previously-observed rhythmicity of collagen fibril formation by fibroblasts, *in vivo* and *in vitro* [10]. Our discovery that the endosomal system is involved in fibril assembly explains how collagen fibrils can be assembled in the absence of endogenous collagen synthesis, namely, if collagen can be retrieved from the extracellular space. We also observed a delay between maximum uptake and maximum fibril numbers. A possible function of this delay may be to increase the concentration of collagen-I in readiness for more efficient fibril initiation. This is supported by our findings that fibroblasts can utilize a solution of Cy3-colI lower than the 0.4 µg/mL critical concentration for fibril formation to assemble fibrils. Of note, the comparison between uptake and fibril numbers are not directly from the same experiments due to experimental limitations; the Cy3/Cy5-colI uptake we observed did not reflect total collagen-I levels, as endogenous collage-I was present as well. Nonetheless, we postulate that the fates of the collagen-I that was taken up by fibroblasts are likely: 1) recycled for fibril formation, 2) re-secreted as protomers, or if they are damaged, 3) degradation. Future research will be required to determine the molecular mechanisms that determines these fates. Surface biotin-labelling in fibroblasts identified two integrin α subunits (α1 and α11) to be absent in VPSko cells which could not make collagen fibrils. There are four known collagen-binding integrins (α1β1, α2β1, α10β1, α11β1), where α2β1 and α11β1 have affinity for fibrillar collagens and are expressed in fibroblasts *in vivo* [23]. α1β1, in contrast, has been demonstrated *in vitro* to display higher affinity for collagen-IV than collagen-I [57]; it has a wide expression pattern *in vivo*, including basement membrane-associated smooth muscle cells, strongly suggesting that collagen-IV is the major ligand for α1β1 integrin instead. Thus, we focused on the effects of integrin α11 subunit. Interestingly, both α1 and α11 subunits were also reduced in VPSoe cells as detected by mass spectrometry; however, validation by western blots indicated that protein levels of integrin α11 follows VPS33B and collagen fibril numbers. We postulate that this discrepancy detected through MS might be due to integrin dynamics at the plasma membrane. Regardless, VPS33B and integrin α11β1 are essential for targeting collagen-I to fibril assembly and not for the secretion of soluble triple helical collagen-I molecules (i.e. protomers); there is also no evidence to support an active role for VPS33B or α11β1 in collagen endocytosis. We have also demonstrated involvement of VPS33B and integrin α11 subunit in IPF, particularly at sites of disease progression (fibroblastic foci). The presence of integrin α11 subunit confirms a recent study utilizing spatial proteomics to decipher regions of IPF tissues, where in addition to integrin α11 the authors also identified proteins involved in collagen biogenesis specific to the fibroblastic foci [58]. Crucially, in our study, levels of *Col1a1* transcript were unchanged between control and IPF fibroblasts, and IHC demonstrated that collagen-I and VPS33B were prevalent in control lungs. This highlights that the production of collagen-I does not equate to fibril formation; indeed, here VPS33B is required for normal collagen homeostasis in the lung, and it is the combination of VPS33B and integrin α11 subunit, coupled with elevated endocytic recycling of exogenous collagen at the fibroblastic focus, that promotes excess collagen-I fibril assemblies, which is a hallmark of fibrosis. Furthermore, this relationship between excessive collagen-I, VPS33B, and integrin α11 is also observed in chronic skin wounds, where, similar to lung fibrosis, there is a chronic unresolved wound-healing response; this is suggestive of a common pathological molecular pathway between the two organs, based on enhanced endocytic recycling of collagen-I protomers directed to fibrillogenesis.

The discovery that VPS33B is required for collagen fibril formation but not collagen protomer secretion highlights an important distinction between ‘collagen secretion’ and ‘collagen fibrillogenesis’, especially in the context of collagen pathologies (e.g. fibrosis, cancer metastasis). In this context, it is crucial to keep in mind that elevated collagen levels (i.e. collagen transcripts or protomer levels) does not equate to the formation of an insoluble fibrillar network. This distinction was also suggested in a recent study on *Pten-*knockout mammary fibroblasts, where SPARC protein acts via fibronectin to affect collagen fibrillogenesis but not secretion [59]. Of note, whilst SPARC and fibronectin were detected in our previously-reported time-series mass spectrometry analyses [10], they were not circadian clock rhythmic in tendon.

The nucleation of a collagen-I fibril is expected to involve collagen-V, a minor fibril-forming collagen that is necessary for the appearance of collagen-I fibrils *in vivo* [60]. Another long-standing view is that fibronectin may tether collagen to a fibronectin-binding integrin and thereby function as a proteolytically cleavable anchor ([61], reviewed by [62]). Whilst the present study did not focus on the role of fibronectin, there is a slight indication that fibronectin is decreased when collagen-I is knocked down, which IF staining suggested that collagen-I and fibronectin do not completely overlap, and that inhibition of endocytosis did not affect fibronectin fibril numbers. Further, biotin-surface labeling mass spectrometry analysis in the present study indicated that the levels of the major fibronectin-binding integrins (integrins α5β1, αVβ3) were not drastically altered between control, VPSko, and VPSoe fibroblasts, indicating that VPS33B controls collagen fibrillogenesis via a fibronectin-independent route. We have recently demonstrated that integrin α11β1 localizes to fibrillar adhesions and contributes to collagen assembly in a mechano-regulated and tensin-1 dependent manner, independent of α5β1 and fibronectin [63]. In these studies, the α11β1 mediated collagen assembly did not appear to depend on endocytosis. In the future it will be important to establish which factors governs endocytosis-dependent and endocytosis-independent α11β1-mediated collagen fibrillogenesis. Interestingly, in fibronectin knockout liver cells, exogenous collagen-V could be added to initiate collagen-I fibrillogenesis [64], which suggests that other matrix proteins could also be recycled in a similar manner as collagen-I. Whilst collagen α1(V) could only be detected in surface biotin-labeled control but not VPSko or VPSoe cells, collagen α2(V) was only detected in VPSoe cells. Of note, we have previously shown that collagen α2(V) protein was rhythmic and in phase with collagen α1(I) and collagen α2(I). Whether collagen-V co-traffics with collagen-I in VPS33B-compartments to fibrillogenic sites at the cell periphery, or collagen-V is directed to the nucleation site by another route, is an important question to address in future studies. Indeed, endocytic recycling of other factors required for collagen fibrillogenesis (e.g. other nucleation factors such as collagen-V, collagen-XI, integrins), in addition to collagen-I protomers, may be required for fibril assembly by fibroblasts.

Previous research has demonstrated that collagen-I can be secreted within minutes [9], whilst other studies have demonstrated a relatively slow emergence of collagen-I fibrils over days in culture [45]. The delay in assembly of fibrils could be due to the requirement to reach the threshold concentration of collagen needed for fibrillogenesis [26]. Procollagen-I can be converted to collagen-I within the secretory pathway [9, 65] in a process that requires giantin for intracellular N-terminal processing of procollagen-I [66]. Thus, it is possible that triple helical collagen-I protomers destined for fibril formation are: 1) directed straight to a VPS33B-positive compartment from the Golgi apparatus, or 2) quickly secreted and then recaptured, via endocytosis, prior to recycling for fibrillogenesis. Such a mechanism would provide the cell with additional opportunities to determine the number of fibrils, the rate at which they are to be initiated, and their required growth in the extracellular matrix (see **Figure 10**). We have not identified the specific proteins that mediate collagen-I uptake into the cell, however as inhibition of clathrin-mediated endocytosis did not completely inhibit collagen endocytosis, it is likely that collagen-I has multiple routes to enter the cells [18–21]. With regards to the fibrillogenesis route, although a recent model of VPS33B/VIPAS39 protein complexes does not consider the possibility that VPS33B can associate directly with vesicular membranes [67], our *in vitro* studies suggest its hydrophobic C-terminal region can act as a tail-anchor domain. Importantly, the proposed model of the VPS33B/VIPAS39 complex suggests the C-terminus of VPS33B would be available for membrane association, consistent with our *in vitro* findings. Further studies will be required to establish whether differences between our models reflect alternative experimental systems, yeast vs. mammalian, and/or the behavior of VPS33B alone vs. in complex with VIPAS39. We found that endogenous VIPAS39 encases collagen-I in intracellular puncta, consistent with a model where VPS33B and VIPAS39 co-traffic with collagen-I at the same endosomal compartment (see also [67]). Our data suggest two membrane bound populations of VPS33B, the majority with a hairpin-like structure and cytosolic C-terminus and a minority that spans the membrane with the C-terminus inside the lumen. Whether or not alternative topologies of VSP33B are associated with different biological functions remains to be determined. Nonetheless, here we have identified a previously unknown role for endocytosis in collagen fibrillogenesis, which is exploited in collagen pathologies.

**Figure 10:**
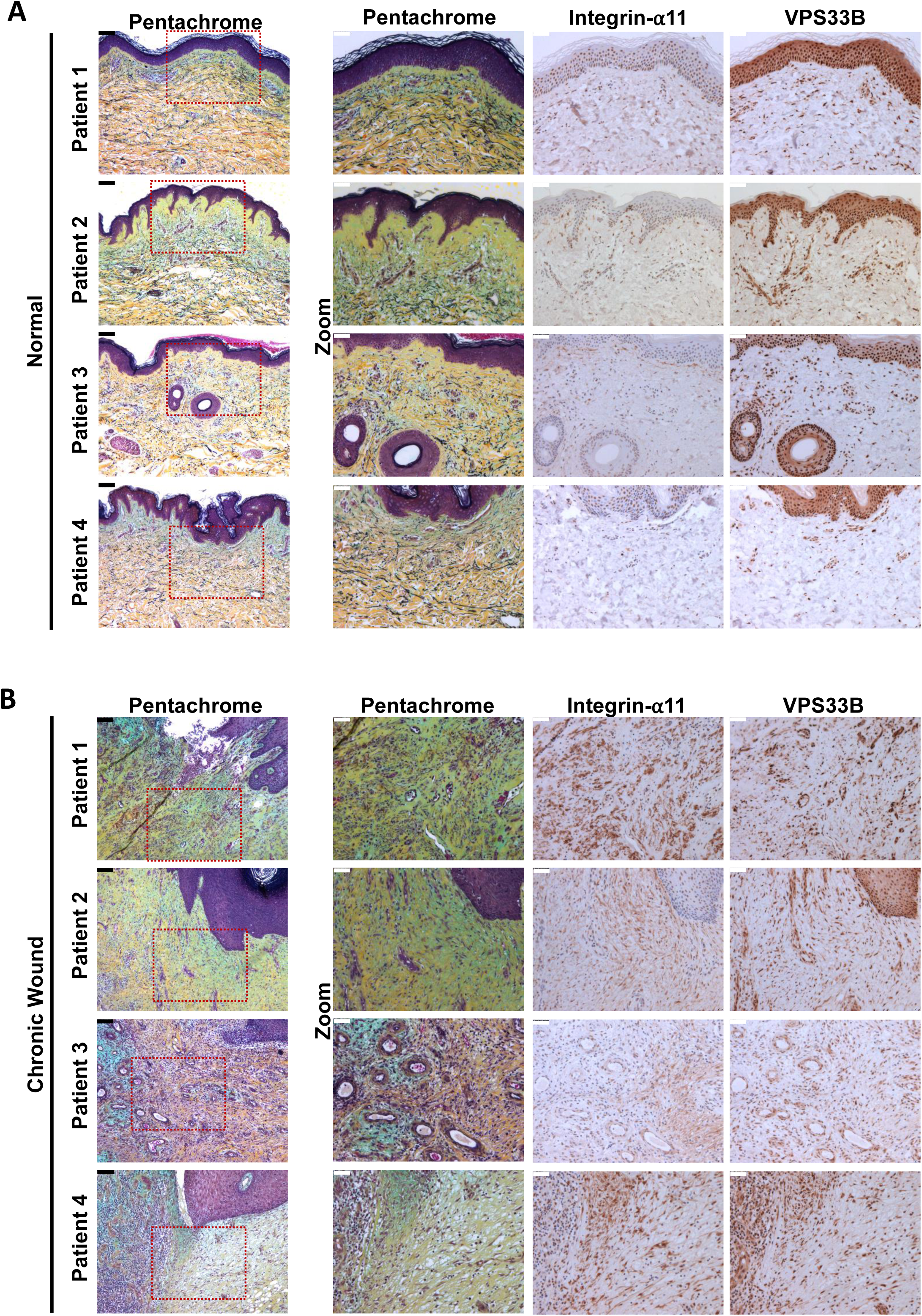
Proteins responsible for collagen fibrillogenesis are also co-localized to diseased areas of chronic skin wounds. A. Immunohistochemistry of 5 µm thick sequential skin sections taken from normal skin regions of patients with chronic skin wounds (Patient 1, Patient 2, Patient 3, Patient 4). Sections were stained with pentachrome, integrin-α11, VPS33B. Scale bars positioned in top left corner: black (unzoomed pentachrome) = 100 µm, white (zoomed sections) = 50 µm. B. Immunohistochemistry of 5 µm thick sequential skin sections taken from the chronic wound areas from patients with chronic skin wounds (Patient 1, Patient 2, Patient 3, Patient 4). Sections were stained with pentachrome, integrin-α11, VPS33B. Scale bars positioned in top left corner: black (unzoomed pentachrome) = 100 µm, white (zoomed sections) = 50 µm.

**Figure 11:**
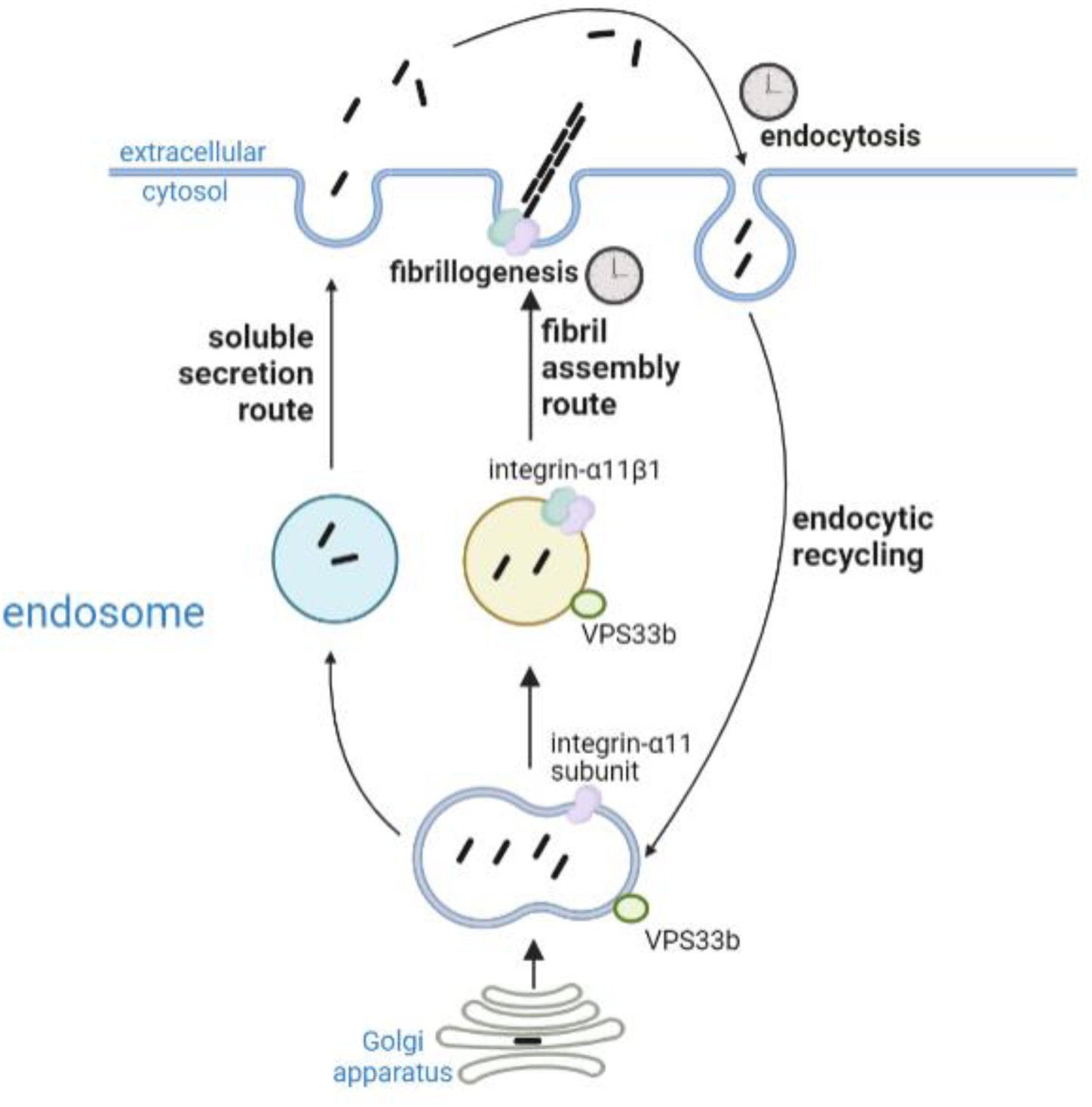
Proposed working model of collagen homeostasis in fibroblasts. Endogenous collagen is either secreted as protomers (soluble secretion route, not circadian rhythmic) or made into fibrils (fibril assembly route, circadian rhythmic). Secreted collagen protomers can be captured by cells through endocytosis (circadian rhythmic) and recycled to make new fibrils. Integrin α11 and VPS33b directs collagen to fibril formation.

## Author contributions

JC, AP and KEK conceived the project. KEK supervised the experiments. AP, JC, RG, JK, TAJ performed experiments and interpreted data. JAH consented and biopsied patients. JAH, LD, MC performed experiments, MH, PC performed super-resolution live imaging, analysis, and data interpretation, YL performed electron microscopy and analysis, CZ produced integrin α11 antibodies and performed antibody screening, DG, RVV, AH provided reagents and data interpretation, and SH and SO’K performed the VPS33B ΔG analysis, topology analyses, and data interpretation. All authors contributed to writing the manuscript.

## Acknowledgements

The proteomics was performed at the Biological Mass Spectrometry Facility with the assistance of Stacey Warwood and Ronan O’Cualain, imaging was performed at the Bioimaging Facility, flow cytometry analyses were performed at the Flow Cytometry Facility with the assistance of Gareth Howell, all in the Faculty of Biology, Medicine and Health (University of Manchester). This report is independent research supported by the North West Lung Centre Charity and National Institute for Health Research Clinical Research Facility at Manchester University NHS Foundation Trust. The views expressed in this publication are those of the author(s) and not necessarily those of the NHS, the North West Lung Centre Charity, National Institute for Health Research or the Department of Health. The authors would like to acknowledge the Manchester Allergy, Respiratory and Thoracic Surgery Biobank and the North West Lung Centre Charity for supporting this project. We would also like to thank the study participants for their contribution. The authors would like to acknowledge and thank Prof David Stephens for his feedback on this manuscript. Some illustrations were created with Biorender.com.

## Materials and Methods

### Procurement of human lung tissue and human lung-derived fibroblasts

The use of human lung tissue was approved by the National Health Research Authority with patient consent (NRES14/NW/0260). The specimens used for this study met the criteria for Idiopathic Pulmonary Fibrosis (IPF) diagnosis [68], as previously described [69]. 4 IPF patient samples and 2 control lung samples were used in this study. The patient-derived fibroblasts, isolated from 5 IPF patient samples and 5 control samples used here were a kind gift from Professor Peter Bitterman and Professor Craig Henke at the University of Minnesota. Briefly, control and IPF-derived fibroblasts (isolated from were established by allowing fibroblasts to propagate from lung tissue explants as previously described [70]). The fibroblasts used in the described experiments were between passages 4 and 6.

### Procurement of human skin tissues

Human chronic wound samples (as defined by [71]) were collected alongside healthy skin samples from 4 consenting patients undergoing surgical debridement and reconstruction at Manchester University NHS Foundation Trust. All procedures were approved by the National Health Research Authority through the ComplexWounds@Manchester Biobank (NRES 18/NW/0847).

### Cell culture

Unless otherwise stated, all cell culture reagents were obtained from Gibco, and all cells were maintained at 37 °C, 5% CO_2_ in a humidified incubator. Immortalized mouse tail tendon fibroblasts (iTTF, isolated from 8-10 weeks old C57BL/6 male mice [10]), and NIH3T3 cells with Dendra2-tagged collagen-I expression (Dendra2-3T3 [45]) were cultured in DMEM/F12 with sodium bicarbonate and L-glutamine (supplemented with 10% fetal calf serum (FCS), 200 μM L-ascorbate-2-phosphate (Sigma), and 10,000 U/mL penicillin/streptomycin), and high-glucose DMEM with sodium bicarbonate and L-glutamine (supplemented with 10% new born calf serum, 200 μM L-ascorbate-2-phosphate, and 10,000 U/mL penicillin/streptomycin) media respectively. Human patient-derived fibroblasts were cultured in low glucose DMEM with sodium bicarbonate and L-glutamine (supplemented with 10% FCS, 200 μM L-ascorbate-2-phosphate, and 10,000 U/mL penicillin/streptomycin). HEK293T cells were cultured in DMEM with sodium bicarbonate and L-glutamine, supplemented with 10% FCS.

To synchronize cells in culture, 100 µM Dexamethasone was added to each sample, then incubated for an hour before the media was removed and replaced with fresh culture media. This marks t=0 in the experiments (i.e. post-synchronisation time 0, or PS0).

Murine tail tendons were isolated by first removing the skin from the mouse tails by cutting it from tail root and degloving the whole tail. Once tendons become exposed, tendon fibers were pulled out using sterile forceps, breaking every other vertebra. The vertebra was then trimmed with scalpels leaving just the tail tendons. These were then placed on a cell culture insert and equilibrated in full DMEM/F12 culture media (see above section) overnight, before the fluorescently-labelled collagen were added at 5μg/ml for 3 days, washed extensively with PBS, and then removed before embedding in low-melting-point agarose for confocal imaging (Leica SP8 upright confocal microscopy). For the pulse-chase experiment, Cy3-labelled collagen was first added and incubated, before tendons were washed and 5FAM-labelled collagen was added for a further 48 hours prior to imaging.

### CRISPR-Cas9-mediated knockout

iTTFs were treated with CRISPR-Cas9 to delete *VPS33B* gene as previously described [10]. Gene knockout was confirmed by western blotting and qPCR.

### Constructs

The cDNA for mouse VPS33B (Uniprot: P59016) was purchased from Genscript (OMu07060D). Sec61β modified with a C-terminal OPG2 tag (residues 1-18 of bovine rhodopsin (Uniprot: P02699) was as previously described [72]. An artificial N-glycosylation site (VPS33B-A606T) and OPG2-tag (VPS33B-OPG2) were incorporated into parent VPS33B by site-directed mutagenesis (Stratagene QuikChange, Agilent Technologies) using the relevant forward and reverse primers (Integrated DNA Technologies; see reagents table) and confirmed by DNA sequencing (GATC, Eurofins Genomics). Linear DNA templates were generated by PCR and mRNA transcribed using T7 polymerase or SP6 RNA polymerase as appropriate (see reagents table). In order to improve the signal intensity of all radiolabeled proteins, an additional five methionine residues (5M) were appended to the C-terminus of all VPS33B linear DNA templates by PCR. All primer combinations used for mutagenesis and PCR are listed in the tables below.

### Transfection and stable infection of over-expression vectors in cells

Three different vectors were used to stably over-express VPS33B protein in iTTF and 3T3 cells – untagged VPS33B overexpression (with red fluorescent protein expression as selection), N-terminus BFP-tagged VPS33B overexpression, and C-terminus BFP-tagged VPS33B overexpression. All proteins were cloned into pLV V5-Luciferase expression vector, where luciferase was excised from the backbone prior to gel purification, or into pCMV expression vector, followed by Gibson Assembly® (NEB) [73]. Lentiviral particles were generated In HEK293T cells, and cells were infected in the presence of 8 μg/mL polybrene (Millipore). Cells were then sorted using Flow Cytometry to isolate RFP-positive or BFP-positive cells. Alternatively, cells were treated with puromycin (Sigma) or G418 (Sigma) depending on selection marker on the vectors.

### Fluorescence labeling of collagen-I

Cy3 or Cy5 NHS-ester dyes (Sigma) were used to fluorescently label rat tail collagen-I (Corning), using a previously described method [25]. Briefly, 3 mg/mL collagen-I gels were made, incubated with 50 mM borate buffer (pH 9) for 15 min, followed with incubation with either Cy3-ester or Cy5-ester (Sigma) dissolved in borate buffer in dark overnight at 4 °C, gently rocking. Dyes were then aspirated and 50 mM Tris (pH 7.5) were added to quench the dye reaction, and incubated in the dark rocking for 10 min. Gels were washed with PBS (with calcium and magnesium ions) 6x, incubating for at least 30 min each wash. The gels were then resolubilized using 500 mM acetic acid and dialyzed in 20 mM acetic acid.

### Circular dichroism of rat tail collagen-I

Circular dichroism spectra of rat tail collagen in 10 mM acetic acid was recorded on a Jasco J810 spectrometer using 0.2mm quartz coverslip sample holders. Spectra were recorded using approximately 0.5 mg/ml rat tail collagen for WT, Cy3 labelled and Cy5 labelled collagens. The melting curves were performed by monitoring at 223 nm only which is the positive triple helical peak and recording between 30 and 70 degrees Celsius, and capturing data points every 30 seconds with a temperature increment of 1 degree every 60 seconds in a 1mm quartz cuvette. All outputs are recorded in machine units rather than being converted to molar units because of the difficulty in ascertaining accurate concentrations of labelled collagen.

### Mass Photometry of rat tail collagen-I

The Refeyn mass photometer was used to assess the mass of the collagen molecules. The instrument was calibrated with BSA and PTX3 with a mass range of 67, 135 and 340 kDa. The WT, Cy3 labelled and Cy5 labelled collagens were diluted to ~20 nM in 10 mM acetic acid and counts read over the course of 60-seconds. The ratiometric contrast was converted to mass using the calibration standards.

### RNA isolation and quantitative real-time PCR

RNA was isolated using TRIzol Reagent (Thermo Fisher Scientific) following manufacturer’s protocol, and concentration was measured using a NanoDrop OneC (Thermo Fisher Scientific). Complementary DNA was synthesized from 1 μg RNA using TaqMan^TM^ Reverse Transcription kit (Applied Biosystems) according to manufacturer’s instructions.

SensiFAST SYBR kit reagents were used in qPCR reactions. Primer sequences can be found in reagents section.

### Protein extraction and western blotting

For lysate experiments, proteins were extracted using urea buffer (8 M urea, 50 mM Tris-HCl pH7.5, supplemented with protease inhibitors and 0.1% β-mercaptoethanol). For conditioned media (CM) media, cells were plated out at 200,000 cells per 6-well plate, and left for 48-72 h before 250 μl was sampled. Samples were mixed with 4xSDS loading buffer with 0.1% β-mercaptoethanol and boiled at 95°C for 5 min. The proteins were separated on either NuPAGE Novex 10% polyacrylamide Bis–Tris gels with 1XNuPAGE MOPS SDS buffer or 6% Tris-glycine gels with 1XTris-glycine running buffer (all Thermo Fisher Scientific), and transferred onto polyvinylidene difluoride (PVDF) membranes (GE Healthcare). The membranes were blocked in 5% skimmed milk powder in PBS containing 0.01% Tween 20. Antibodies were diluted in 2.5% skimmed milk powder in PBS containing 0.01% Tween 20. The primary antibodies used were: rabbit polyclonal antibody (pAb) to collagen-I (1:1,000; Gentaur), mouse mAb to vinculin (1:1000; Millipore), rabbit pAb to integrin α11 subunit (1:1000; see [74]) and mouse mAb to VPS33B (1:500; Proteintech). Horseradish-peroxidase-conjugated antibodies and Pierce ECL western blotting substrate (both from Thermo Fisher Scientific) were used and reactivity was detected on GelDoc imager (Biorad). Alternatively, Licor goat-anti-mouse 680, mouse-anti-rabbit 800 were used and reactivity detected on an Odyssey Clx imager.

### *In vitro* ER import assays

Translation and ER import assays were performed in nuclease-treated rabbit reticulocyte lysate (Promega) as previously described [38, 39, 75]: briefly, in the presence of EasyTag EXPRESS ^35^S Protein Labelling Mix containing [^35^S] methionine (Perkin Elmer) (0.533 MBq; 30.15 TBq/mmol), 25 μM amino acids minus methionine (Promega), 6.5% (v/v) nuclease-treated ER-derived canine pancreatic microsomes (from stock with OD_280_=44/mL) or an equivalent volume of water, and 10% (v/v) of *in vitro* transcribed mRNA (~1 μg/μL) encoding relevant precursor proteins. Translation reactions for Sec61βOPG2 (20 μL; 1X) were performed at 30°C for 30 min whereas VPS33B and its variants (20 μL; 2X) were performed at 30°C for 1 h. Irrespective of the precursor protein, all translation reactions were finished by incubating with 0.1 mM puromycin for 10 min at 30°C to ensure translation termination and the ribosomal release of newly synthesized proteins prior to analysis. Microsomal membrane-associated fractions were recovered by centrifugation through an 80 μL high-salt cushion (0.75 M sucrose, 0.5 M KOAc, 5 mM Mg(OAc)_2_ and 50 mM HEPES-KOH, pH 7.9) at 100,000 g for 10 min at 4°C, the pellet suspended directly in SDS sample buffer and, where indicated, treated with 1000 U of endoglycosidase Hf (New England Biolabs, P0703S). All samples were denatured for 10 min at 70°C and resolved by SDS–PAGE (10% or 16% PAGE, 120 V, 120-150 min). Gels were fixed for 5 min (20% MeOH, 10% AcOH), dried for 2 h at 65°C, and radiolabeled products were visualized using a Typhoon FLA-700 (GE Healthcare) following exposure to a phosphorimaging plate for 24–72 h. Images were opened using Adobe Photoshop and annotated using Adobe Illustrator.

### Flow cytometry and imaging

Cy3- or Cy5-tagged collagen was added to cells and incubated at 37°C, 5% CO_2_ in a humidified incubator for predetermined lengths of times before washing in PBS, trypsinized, spun down at 2500 rpm for 3 min at 4 °C, and resuspended in PBS on ice. For labelled dextran experiment, 200 μg/ml Oregon green^TM^ 488-labelled 70kDa dextran and 10 μg/ml Cy5-colI were added to the cells together and incubated for 1 hr; for unlabelled collagen-I saturation experiments, 100 μg/ml rat-tail collagen-I were added together with 10 μg/ml labelled collagen-I and incubated for 1 hr. Cells were then processed as described above. Cells were analyzed for Cy3-/Cy5-collagen/488-dextran uptake using LSRFortessa (BD Biosciences). For imaging, cells were prepared as described, and analyzed on Amnis^®^ ImageStream^®X^Mk II (Luminex).

### Decellularization of cells to obtain extracellular matrix

Cells were seeded out at 50,000 in a 6-well plate and cultured for 7 days before decellularization. Extraction buffer (20mM NH_4_OH, 0.5% Triton X-100 in PBS) was gently added to cells and incubated for 2 minutes. Lysates were aspirated and the matrix remaining in the dish were washed gently twice with PBS, before being scraped off into ddH_2_O for further processing.

### Hydroxyproline assay

Samples were incubated overnight in 6 M HCl (diluted in ddH_2_O (Fluka); approximately 1 mL per 20 mg of sample) in screw-top tubes (StarLab) in a sand-filled heating block at 100 °C covered with aluminum foil. The tubes were then allowed to cool down and centrifuged at 12,000 g for 3 min. Hydroxyproline standards were prepared (starting at 0.25 mg/mL; diluted in ddH2O) and serially diluted with 6 M HCl. Each sample and standard (50 μL) were transferred into fresh Eppendorf tubes, and 450 μL chloramine T reagent [0.127 g chloramine T in 50% N-propanol diluted with ddH2O; made up to 10 mL with acetate citrate buffer (120 g sodium acetate trihydrate, 46 g citric acid, 12 mL glacial acetic acid, 34 g sodium hydroxide) adjusted to pH 6.5 and then made to 1 litre with dH_2_O; all reagents from Sigma] was added to each tube and incubated at room temperature for 25 min. Ehrlich’s reagent (500 μL; 1.5 g 4-dimethylaminobenzaldehyde diluted in 10 mL N-propanol:perchloric acid (2:1)) was added to each reaction tube and incubated at 65 °C for 10 min and then the absorbance at 558 nm was measured for 100 μL of each sample in a 96-well plate format.

### Electron microscopy

Unless otherwise stated, incubation and washes after EM fixation were done at room temperature. Cells were plated on top of ACLAR films and allows to deposit matrix for 7 days. The ACLAR was then fixed in 2% glutaraldehyde/100mM phosphate buffer (pH 7.2) for at least 2 h and washed in ddH_2_O 3 x 5 min. The samples were then transferred to 2% osmium (v/v)/1.5% potassium ferrocyanide (w/v) in cacodylate buffer (100 mM, pH 7.2) and further fixed for 1 h, followed by extensive washing in ddH_2_O. This was followed by 40 minutes of incubation in 1% tannic acid (w/v) in 100 mM cacodylate buffer, and then extensive washes in ddH_2_O. and placed in 2% osmium tetroxide for 40 min. This was followed by extensive washes in ddH_2_O. Samples were incubated with 1% uranyl acetate (aqueous) at 4 °C for at least 16 h, and then washed again in ddH_2_O. Samples were then dehydrated in graded ethanol in the following regime: 30%, 50%, 70%, 90% (all v/v in ddH_2_O) for 8 min at each step. Samples were then washed 4 x 8 min each in 100% ethanol, and transferred to pure acetone for 10 min. The samples were the infiltrated in graded series of Agar100Hard in acetone (all v/v) in the following regime: 30% for 1 h, 50% for 1 h, 75% for overnight (16 h), 100% for 5 h. Samples were then transferred to fresh 100% Agar100Hard in labeled moulds and allowed to cure at 60 °C for 72 h. Sections (80 nm) were cut and examined using a Tecnai 12 BioTwin electron microscope.

### Surface biotinylation-pulldown mass spectrometry

Cells were grown in 6 well plates for 72 h prior to biotinylating. Briefly, cells around 90% confluence were kept on ice and washed in ice-cold PBS, following with incubation with ice-cold biotinylating reagent (prepared fresh, 200 µg/mL in PBS, pH 7.8) for 30 min at 4 °C gently shaking. Cells were then washed twice in ice-cold TBS (50 mM Tris, 100 mM NaCl, pH 7.5), and incubated in TBS for 10 min at 4 °C. Cells were then lysed in ice-cold 1% TritonX (in PBS, with protease inhibitors) and lysates cleared by centrifugation at 13,000 g for 10 min at 4 °C. Supernatant was then transferred to a fresh tube and 1/5 of the lysates were kept as a reference sample. Streptavidin-Sepharose beads was aliquoted into each sample and incubated for 30 min at 4 °C rotating. Beads were then washed 3x in ice cold PBS supplemented with 1% TritonX, followed by one final wash in ice cold PBS and boiled at 95 °C for 10 min in 2x sample loading buffer, followed by centrifugation at 13,000 g for 5 min follow by sample preparation for Mass Spectrometry analysis. The protocol used for sample preparation was as described previously [69]. Mass spectrometry results files were exported into Proteome Discoverer (PD) for identification and spectral counting. All searches included the fixed modification for carbamidomethylation on cysteine residues resulting from IAA treatment to prevent cysteine bonding. The variable modifications included in the search were oxidized methionine (monoisotopic mass change, +15.955 Da); hydroxylation of asparagine, aspartic acid, proline, or lysine (monoisotopic mass change, +15.955 Da); and phosphorylation of threonine, serine, and tyrosine (79.966 Da). A maximum of 2 missed cleavages per peptide was allowed. The minimum precursor mass was set to 350Da with a maximum of 5000. Precursor mass tolerance was set to 10ppm, fragment mass tolerance was 0.02Da and minimum peptide length was 6. Peptides were searched against the Swissprot database using Sequest HT with a maximum false discovery rate of 1%. Proteins required a minimum FDR of 1% and were filtered to remove known contaminants and to have at least 2 unique peptides. Missing values were assumed to be due to low abundance. For spectral counting, enriched samples were normalized and spectral matches were compared. Peptide spectral matches were only included if they were calculated to have a q-value of less than 0.01.

### Imaging and immunofluorescence

For fluorescence live imaging of Cy3-colI internalisation, iTTF cells were seeded for 24 h onto Ibidi µ-plates and stained using CellMask™ Green Plasma Membrane Stain (Invitrogen C37608) according to the manufacturers protocol. Cy3-colI was added to cells and images were collected in a 37**°**C chamber using a Zeiss 3i spinning disk (CSU-X1, Yokagowa) confocal microscope with a 63x/1.40 Plan-Apochromat oil objective. 7.5 µm z-stacks with a step size of 0.5 µm were captured at 2 minute intervals using 488 nm (100 power, 150 ms exposure) and 561 nm (100 power, 100 ms exposure) lasers with SlideBook 6.0 software (3i) and a front illuminated Prime sCMOS camera. Imaris 9.9.1 software was then used to generate 3D reconstructions.

For fixed immunofluorescence imaging, cells plated on coverslips or Ibidi μ-plates were fixed with 100% methanol at −20°C and then permeabilized with 0.2% Triton-X in PBS. Primary antibodies used were as follows: rabbit pAb collagen-I (1:400, Gentaur OARA02579), rabbit pAb VIPAS (1:50, Proteintech 20771-1-AP), mouse mAb FN1 (1:400, Sigma F6140 or Abcam ab6328). Secondary antibodies conjugated to Alexa-488, Alexa-647, Cy3, and Cy5 were used (ThermoScientific), and nuclei were counterstained with DAPI (Sigma). Coverslips were mounted using Fluoromount G (Southern Biotech). Images were collected using a Leica SP8 inverted confocal microscope (Leica) using an x63/0.50 Plan Fluotar objective. The confocal settings were as follows: pinhole, 1 Airy unit; scan speed, 400 Hz bidirectional; format 1024 x 1024 or 512 x 512. Images were collected using photomultiplier tube detectors with the following detection mirror settings: DAPI, 410-483; Alexa-488, 498-545nm; Cy3, 565-623 nm; Cy5, 638-750 nm using the 405 nm, 488 nm, 540 nm, 640 nm laser lines. Alternatively, images were collected using EVOS^TM^ M7000 (Thermo Fisher) using 40x/1.3na oil objective. Images were collected in a sequential manner to minimize bleed-through between channels.

For fluorescence live-imaging, cells were plated onto Ibidi μ-plates and imaged using Zeiss LSM880 NLO (Zeiss). For split-GFP experiment, cells were seeded for 24 h onto Ibidi μ-plates before imaging with an Olympus IXplore SpinSR (Olympus) with 100x oil magnification. Prior to imaging, media was changed to FluoroBrite media with the appropriate supplements.

### Immunohistochemistry

Human lung samples were formalin-fixed and paraffin-embedded (FFPE). 5-μm FFPE sections were obtained and mounted onto Superfrost® Plus (Thermo Scientific™) slides and subjected to antigen heat retrieval using citrate buffer (Abcam, ab208572), in a pre-heated steam bath for 20 minutes, before cooling to room temperature in a water bath for 20 min. Slides were then treated with 3-4% hydrogen peroxide (Leica Biosystems RE7101) for 10 min, blocked in SuperBlock™ (TBS) blocking buffer for a minimum of 1 h (Thermo Scientific™; 37581), and probed with primary antibodies overnight at 4°C in 10% SuperBlock™ solution in Tris Buffered Saline Tween 20 solution (TBS-T, pH 7.6). Primary antibodies used were as follows: Collagen-I A1/A2 (Rockland Immunochemicals Inc 600-401-103.0.5), integrin α11 (integrin α11 mAb 210F4 [76]), VPS33B (Atlas antibodies).

After overnight incubation, specimens were subjected to Novolink Polymer Detection Systems (Leica Biosystems RE7270-RE, as per the manufacturer’s recommendations), with multiple TBS-T washes. Sections were developed for 5 min with DAB Chromagen (Cell Signaling Technology®, 11724/5) before being counterstained with haematoxylin for 1 minute, followed by acid alcohol and blueing solution application. Slides were dehydrated through sequential ethanol and xylene before being cover-slipped with Permount mounting medium (Thermo Scientific™, SP15).

### Histological imaging

Stained slides were imaged using a DMC2900 Leica camera along with Lecia Application Suite X software (Leica).

## QUANTIFICATION AND STATISTICAL ANALYSIS

Data are presented as the mean ± s.e.m. unless otherwise indicated in the figure legends. The sample number ‘N’ indicates the number of independent biological samples in each experiment, and ‘n’ indicates the number of technical repeats, and are indicated in the figure legends. Data were analyzed as described in the legends. The data analysis was not blinded, apart from quantification of fibril numbers over time, and differences were considered statistically significant at *P*<0.05, using Student’s t-test or One-way Anova, unless otherwise stated in the figure legends. Analyses were performed using Graphpad Prism 8 or 10.2.0 software. Significance levels are: * *P*<0.05; ** *P*<0.01; *** *P*<0.005; \*\*\*\**P*<0.0001. Where applicable, normality test was performed using the Shapiro-Wilk method. For periodicity, analysis was performed using the MetaCycle package[77] in the R computing environment [78] with the default parameters.

### Reagents

| REAGENT or RESOURCE | SOURCE | IDENTIFIER |
| --- | --- | --- |

| Antibodies |  |  |
| --- | --- | --- |
| rabbit polyclonal antibody (pAb) to collagen-I | Aviva Systems Biology | Cat# OARA02579, RRID:AB_10873334 |
| rabbit polyclonal antibody (pAb) to collagen-I | Kerafast | Cat# ENH018 |
| mouse mAb to mouse vinculin | Sigma-Aldrich | Cat# V9131, RRID:AB_477629 |
| rabbit pAb to mouse integrin α11 subunit | This paper | N/A |
| mouse mAb to mouse VPS33B | Proteintech | Cat# 12195-1-AP, RRID:AB_2215198 |
| Rabbit pAb to mouse VIPAS | Proteintech | Cat# 20771-1-AP, RRID:AB_10695764 |
| Mouse mAb to FN1 | Sigma-Aldrich | Cat# F6140, RRID:AB_476981 |
| Mouse mAb to FN1 | Abcam | Cat# ab6328, RRID:AB_305428 |
| Rabbit pAb to human VPS33B | Atlas antibodies | Cat# HPA040415, RRID:AB_10795419 |
| Mouse mAb integrin α11 210F4 | DOI: 10.1002/cjp2.148 | N/A |
| Rabbit pAb to Collagen-I | Rockland Immunochemicals | 600-401-103.0.5, RRID:AB_217595 |
| IRDye® 680RD Goat anti-Mouse IgG (H + L) | Li-Cor | Cat# 925-68070, RRID:AB_2651128 |
| IRDye® 800RD Goat anti-Rabbit IgG (H + L) | Li-Cor | Cat# 925-32211, RRID:AB_2651127 |
| Goat anti-Mouse IgG (H+L) Secondary Antibody, HRP | Thermo Fisher Scientific | Cat# 32430, RRID:AB_1185566 |
| Goat anti-Rabbit IgG (H+L) Secondary Antibody, HRP | Thermo Fisher Scientific | Cat# 32460, RRID:AB_1185567 |
| Goat anti-Rabbit IgG (H+L) Cross-Adsorbed Secondary Antibody, Cyanine3 | Thermo Fisher Scientific | Cat# A10520, RRID:AB_2534029 |
| Goat anti-Rabbit IgG (H+L) Cross-Adsorbed Secondary Antibody, Cyanine5 | Thermo Fisher Scientific | Cat# A10523, RRID:AB_2534032 |
| Goat anti-Mouse IgG (H+L) Cross-Adsorbed Secondary Antibody, Cyanine3 | Thermo Fisher Scientific | Cat# A10521, RRID:AB_2534030 |
| Goat anti-Mouse IgG (H+L) Highly Cross-Adsorbed Secondary Antibody, Alexa Fluor 647 | Thermo Fisher Scientific | Cat# A-21236, RRID:AB_2535805 |
| Goat anti-Mouse IgG (H+L) Highly Cross-Adsorbed Secondary Antibody, Alexa Fluor 488 | Thermo Fisher Scientific | Cat# A-11029, RRID:AB_2534088 |
| Goat anti-Rabbit IgG (H+L) Highly Cross-Adsorbed Secondary Antibody, Alexa Fluor 488 | Thermo Fisher Scientific | Cat# A-11034, RRID:AB_2576217 |
| Bacterial and virus strains |  |  |
| pLV-V5-VPS33B-GFP1-10 | This paper | N/A |
| pLV-V5-VPS33B-BFP | This paper | N/A |
| pLV-V5-BFP-VPS33B | This paper | N/A |
| pCMV-SBP-GFP11-COL1A1 | This paper | N/A |
| pcDNA3.1+/C-(K)-DYK | This paper | N/A |
| Biological samples |  |  |
| Human control lung tissues | Manchester University NHS Foundation Trust | NRES14/NW/0260 |
| Human IPF lung tissues | Manchester University<br>NHS Foundation Trust | NRES14/NW/0260 |
| Human chronic skin wound tissue | Manchester University<br>NHS Foundation Trust | NRES 18/NW/0847 |
| Human healthy skin wound tissue | Manchester University<br>NHS Foundation Trust | NRES 18/NW/0847 |
| Chemicals, peptides, and recombinant proteins |  |  |
| Recombinant *S. pyogenes* Cas9 nuclease protein | IDT | 1081059 |
| Rat tail collagen-I | Corning | 354259 |
| Dyngo®4a | Abcam | Ab120689 |
| TRIzol reagent | Invitrogen | 15596-018 |
| Oregon green™ 488-Dextran, 70kDa | Life technologies | D7173 |
| Cy®3 NHS Ester | Sigma | GEPA13101 |
| Cy®5 NHS Ester | Sigma | GEPA15101 |
| Critical commercial assays |  |  |
| Gibson Assembly Cloning kit | NEB | E5510S |
| Site-Directed Mutagenesis QuikChange kit | Agilent | 200513 |
| Novolink Polymer Detection Systems | Leica Biosystems | RE7270-RE |
| Rabbit reticulocyte lysate, nuclease treated | Promega | L4960 |
| TaqMan™ Reverse Transcription kit | Applied Biosystems | N8080234 |
| SensiFAST™SYBR® No-ROX kit | Bioline | BIO-98005 |
| Experimental models: Cell lines |  |  |
| NIH3T3-Dendra2-Coll | This paper | N/A |
| Immortalized mouse tail tendon fibroblasts | This paper | N/A |
| Primary human IPF lung fibroblasts | University of<br>Minnesota | N/A |
| Primary human control lung fibroblasts | University of<br>Minnesota | N/A |
| HEK293T | This paper | N/A |
| Oligonucleotides |  |  |
| siCol1a1, Mission® esiRNA | Merck | EMU069551 |
| siltga11, Mission® esiRNA | Merck | EMU042761 |
| Primers for site-directed mutagenesis | See primer list | N/A |
| Primers for transcription templates | See primer list | N/A |
| Primers for qPCR reactions | See primer list | N/A |
| Recombinant DNA |  |  |
| VPS33B mouse cDNA | Genscript | Omu07060D |
| Sec61β modified with a C-terminal OPG2 tag | This paper | N/A |
| VPS33B A-606-T | This paper | N/A |
| VPS33B-OPG2 | This paper | N/A |
| VPS33B-Sec61βTMD | This paper | N/A |
| VPS33B-Sec61βTMD-OPG2 | This paper | N/A |
| Software and algorithms |  |  |
| Fiji ImageJ | doi:10.1038/nmeth.2019 | https://imagej.net/software/fiji/ |
| MetaCycle | doi:10.1093/bioinformatics/btw405 | https://github.com/gangwug/MetaCycle |
| Scaffold Proteome Software | doi:10.1002/pmic.200900437 | https://www.proteomesoftware.com/products |
| Graphpad Prism 8 | https://www.graphpad.com/scientific-software/prism/ | https://www.graphpad.com/scientific-software/prism/ |
| Biorender | https://biorender.com/ | https://biorender.com/ |

Primer list for cDNAs generated by site-directed mutagenesis.

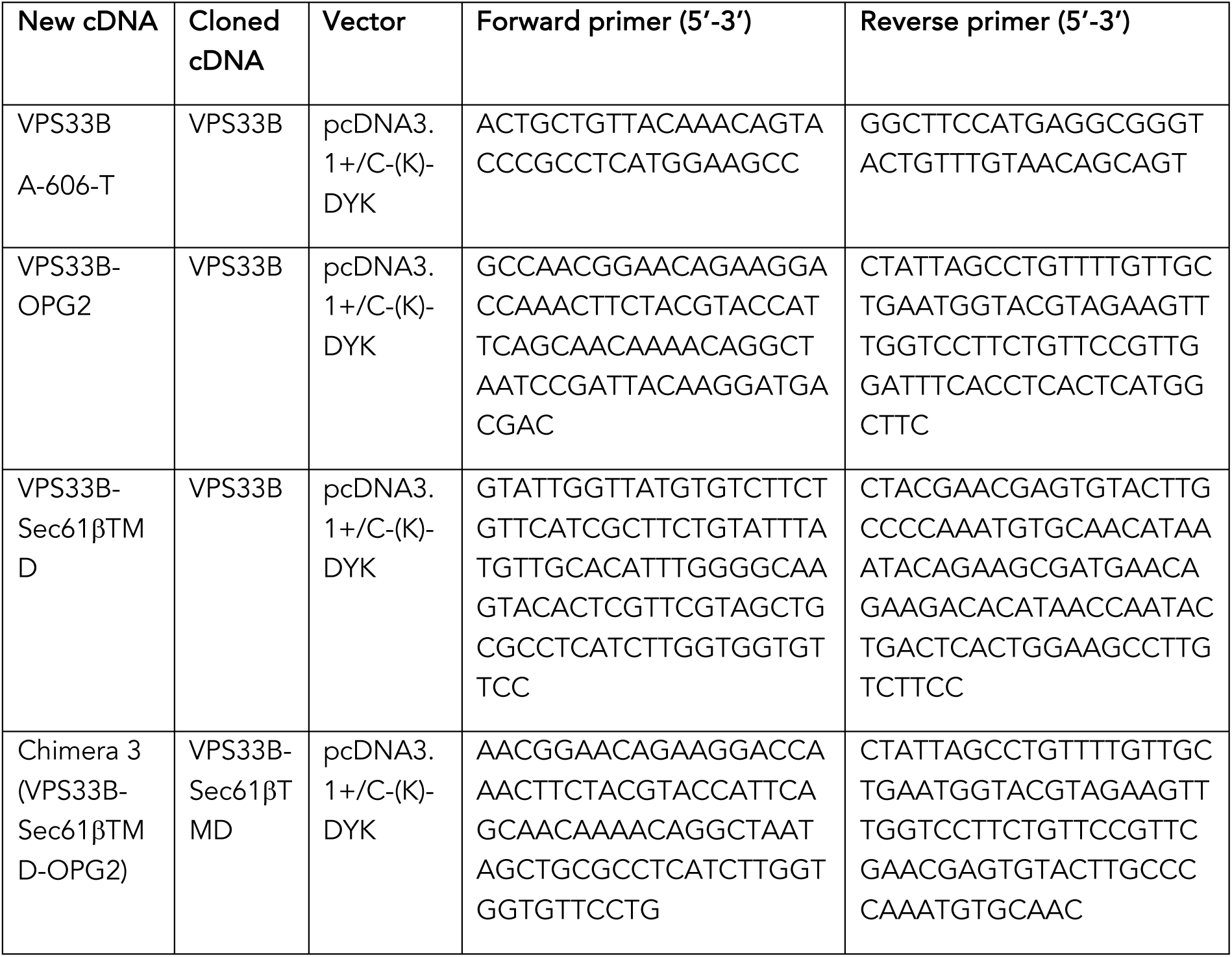

Primer list for PCR reactions used to create transcription templates.

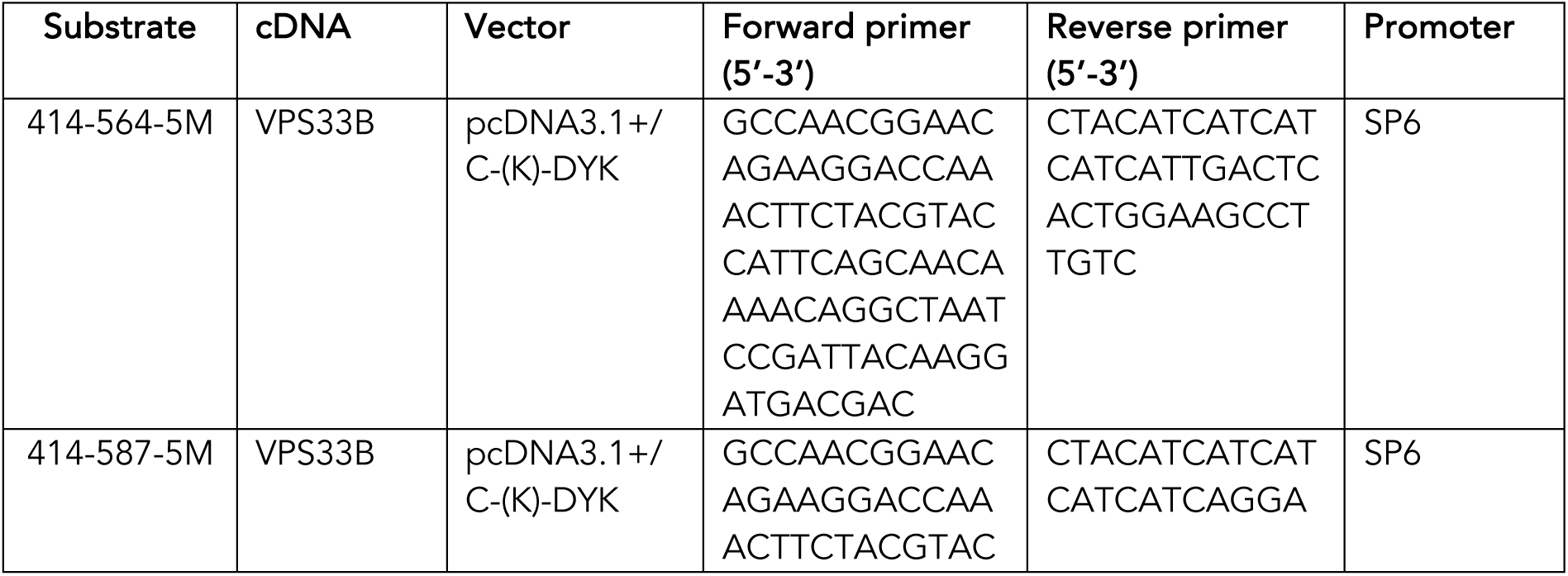
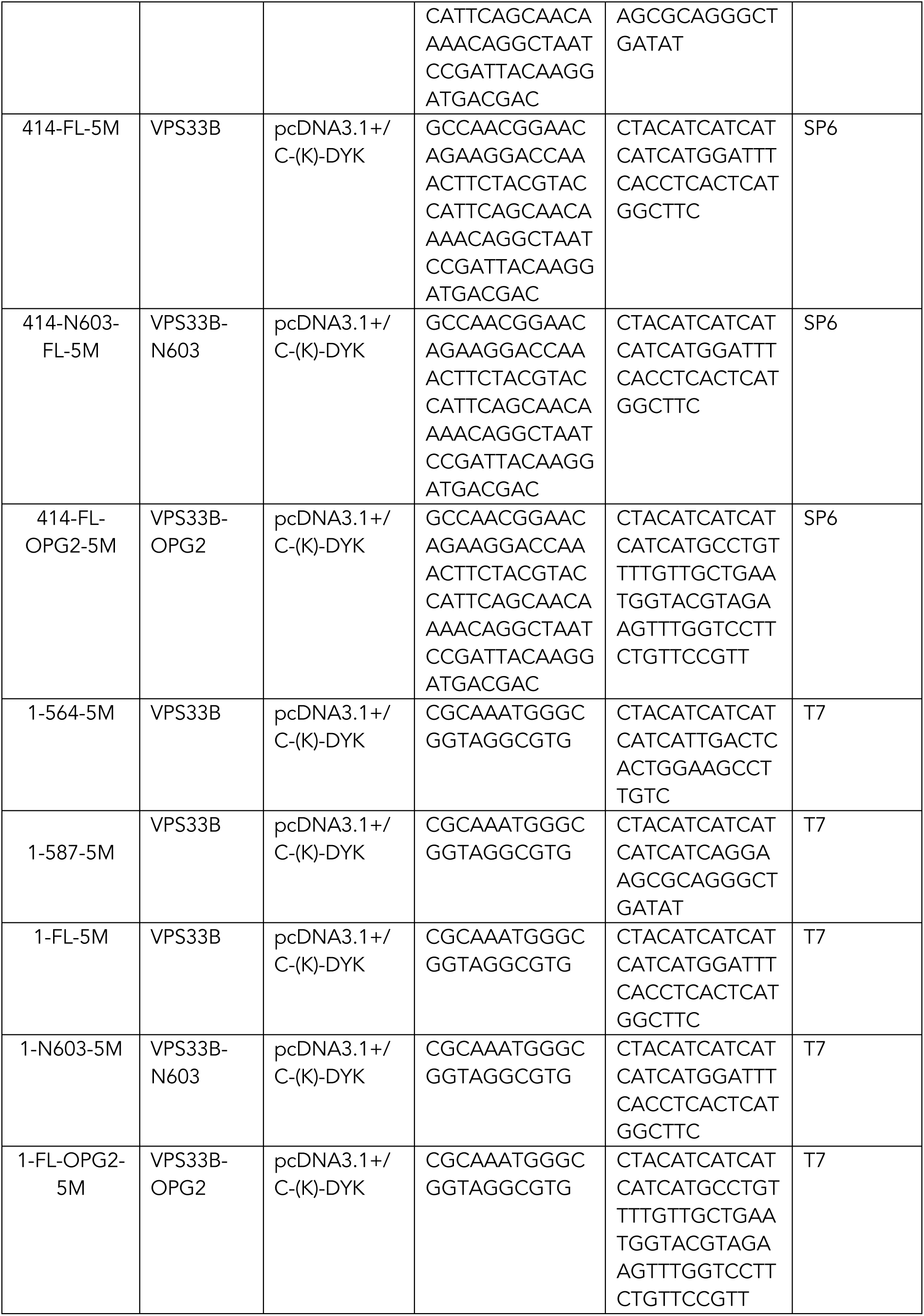

Primer list for qPCR reactions.

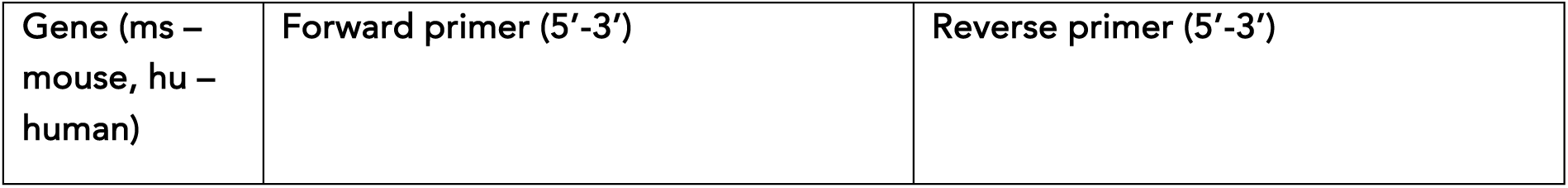
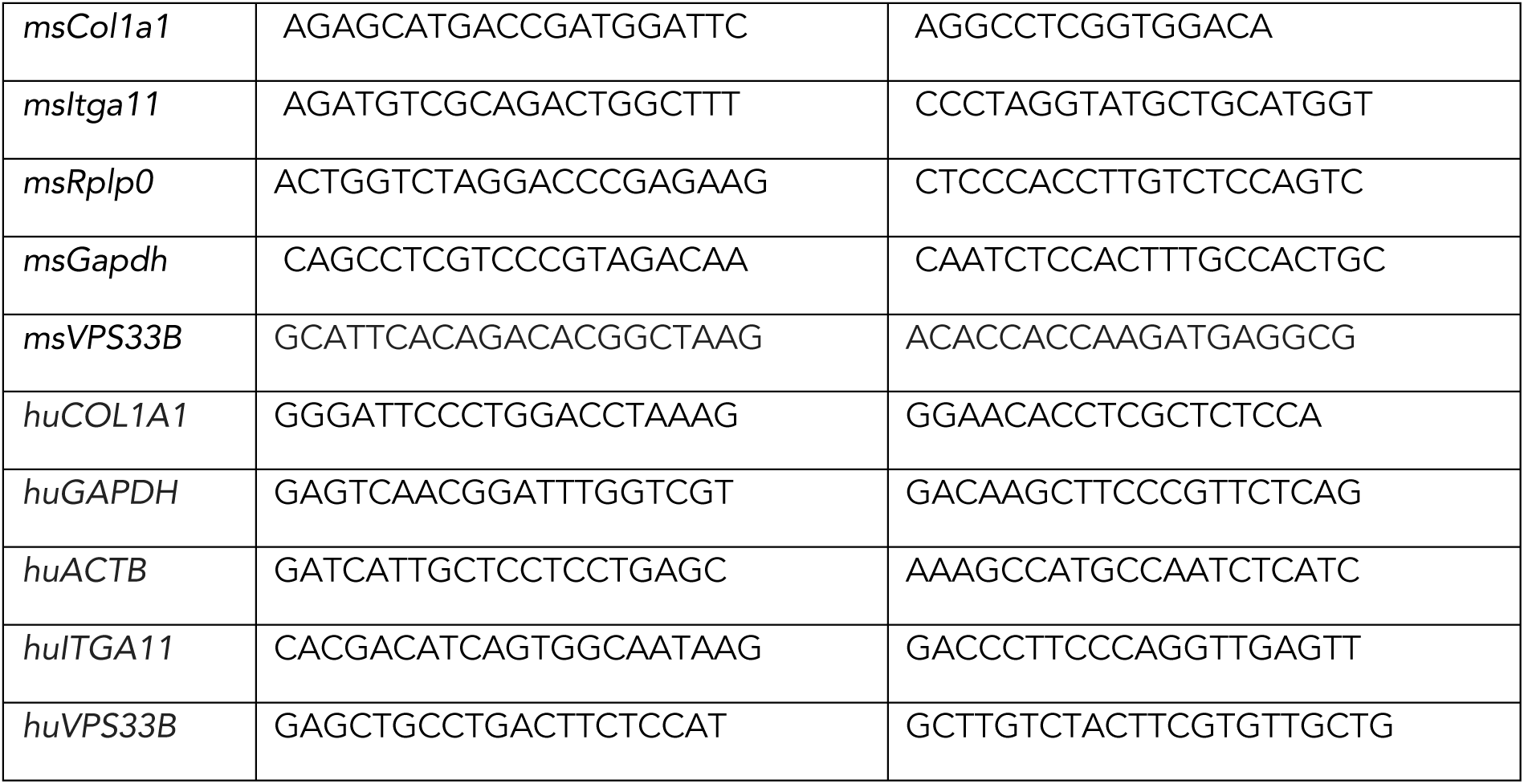

