## Supplementary figures for "Endocytic recycling is central to circadian collagen fibrillogenesis and disrupted in fibrosis"

#### Figure S1

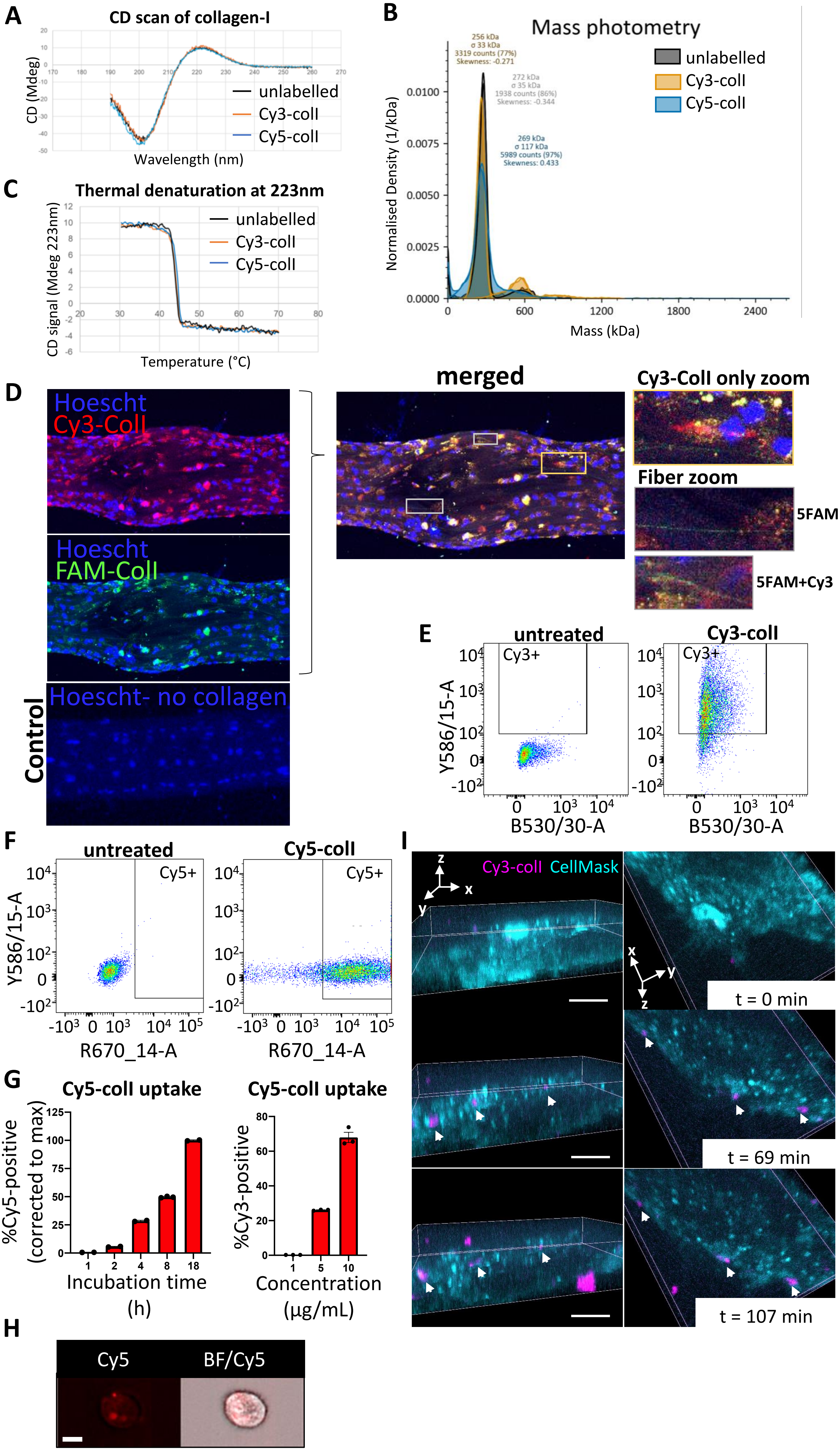

A
CD scan of collagen-I
CD (Mdeg)
Wavelength (nm)
unlabelled
Cy3-coll
Cy5-coll
B
Mass photometry
unlabelled
Cy3-coll
Cy5-coll
256 kDa
σ 33 kDa
3319 counts (77%)
Skewness: -0.271
272 kDa
σ 35 kDa
1938 counts (86%)
Skewness: -0.344
269 kDa
σ 117 kDa
5989 counts (97%)
Skewness: 0.433
Normalised Density (1/kDa)
Mass (kDa)
C
Thermal denaturation at 223nm
CD signal (Mdeg 223nm)
Temperature (°C)
unlabelled
Cy3-coll
Cy5-coll
D
Hoescht
Cy3-Coll
Hoescht
FAM-Coll
Control
Hoescht- no collagen
merged
Cy3-Coll only zoom
Fiber zoom
5FAM
5FAM+Cy3
E
untreated
Cy3-coll
Cy3+
Y586/15-A
B530/30-A
F
untreated
Cy5-coll
Cy5+
Y586/15-A
R670_14-A
G
Cy5-coll uptake
%Cy5-positive (corrected to max)
Incubation time (h)
Cy5-coll uptake
%Cy3-positive
Concentration (µg/mL)
H
Cy5
BF/Cy5
I
Cy3-coll
CellMask
t = 0 min
t = 69 min
t = 107 min

**Figure S1**

A. Circular dichroism spectra of unlabelled (black), Cy3-labelled (orange, Cy3-colI) and Cy5-labelled (blue, Cy5-colI) collagen-I in acetic acid showing the helical positive peak at 223 nm in each of the spectra.

B. Mass photometry of unlabelled (black), Cy3-labelled (orange, Cy3-colI) and Cy5-labelled (blue, Cy5-colI) collagen-I in acetic acid.

C. Temperature induced thermal unfolding of collagen-I monitored at 223 nm. The thermal transition is the same for each of the labelled and non-labelled collagen-I with a mi-point melting temperature of 44 degrees Celsius.

D. Fluorescent images of tail tendon either not incubated with fluorescently-labeled collagen-I (Control, bottom), or incubated with fluorescently-labeled collagen-I for 5 days – Cy3-colI as added for the first 3 days, removed, and then with 5FAM-labeled collagen-I (FAM-colI) added in the last two days. Images show presence of collagen-I within the cells, and fibril-like fluorescence signals outside of cells. Hoechst stain was used to locate cells within the tendon. Area surrounded by yellow box expanded on the right highlighting a cell with only Cy3-colI present intracellularly. Area surrounded by grey boxes expanded on the right, highlighting fibril-like fluorescence signals that are either FAM-colI positive only, or have co-localization of both Cy3-colI and FAM-colI. Representative of N=2.

E. Representative dot plots from flow cytometry analysis representing Cy3 gates used in control and iTTFs incubated with 1 µg/mL Cy3-labeled collagen (Cy3-colI) for 18 h. Representative of N>3.

F. Representative dot plots from flow cytometry analysis representing Cy5 gates used in control and iTTFs incubated with 1 µg/mL Cy5-labeled collagen (Cy5-colI) for 18 hours. Representative of N>3.

G. Bar chart showing a progressive increase of percentage of fluorescent iTTFs incubated with 1.5 µg/mL Cy5-colI over time (left), and an increase of percentage fluorescent iTTFs incubated with increasing concentration of Cy5-colI for one hour (right), suggesting a non-linear time-dependent and dose-dependent uptake pattern. N=3.

H. Flow cytometry imaging of iTTFs incubated with 5 µg/mL Cy5-labeled collagen-I for one hour, showing that collagen-I is taken up by cells and held in vesicular structures. Images acquired using ImageStream at 40x magnification. Scale bar = 10 $\mu$m. Cy5 – Cy5 channel, BF/Cy5 – merged image of BF and Cy5. Representative of >500 cells images collected per condition.

I. Fluorescent image series of iTTFs incubated with Cy3-labeled collagen-I at 0 min (t = 0min) 69 min (t = 69 min) and 107 min (t = 107 min) after addition (rows) and viewed from different angles (columns), with cell mask (cyan) to distinguish the cell volume. White arrows point to intracellular collagen in vesicular like structures. Representative of N = 3. Scale bar = 20 µm.

**Figure S2**

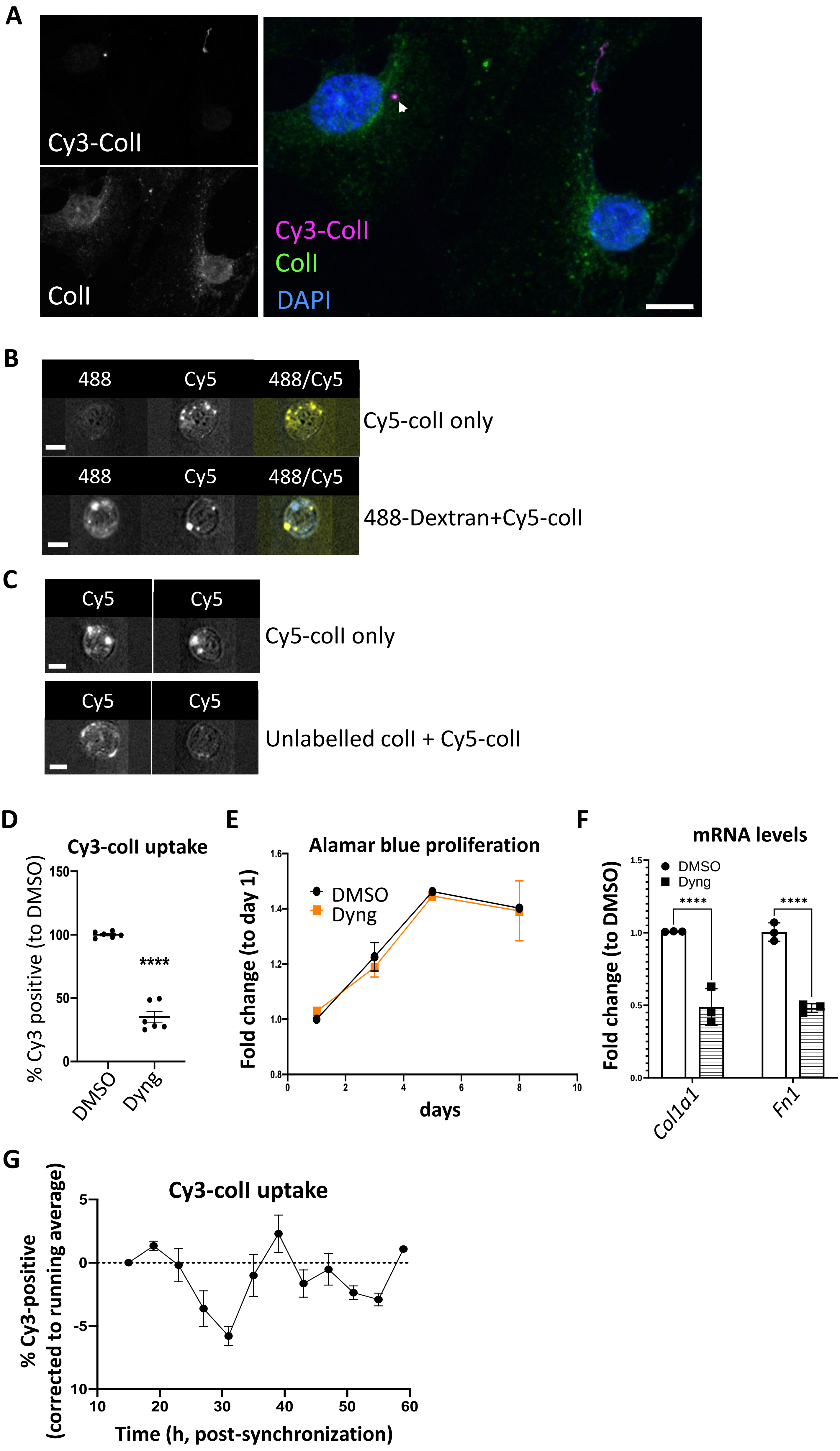
A
Cy3-CollI
CollI
Cy3-CollI
CollI
DAPI
B
488
Cy5
488/Cy5
Cy5-collI only
488
Cy5
488/Cy5
488-Dextran+Cy5-collI
C
Cy5
Cy5
Cy5-collI only
Cy5
Cy5
Unlabelled collI + Cy5-collI
D
Cy3-collI uptake
% Cy3 positive (to DMSO)
150
100
50
0
****
DMSO
Dyng
E
Alamar blue proliferation
Fold change (to day 1)
1.6
1.4
1.2
1.0
0.8
DMSO
Dyng
0
2
4
6
8
10
days
F
mRNA levels
Fold change (to DMSO)
1.5
1.0
0.5
0.0
DMSO
Dyng
****
****
Col1a1
Fn1
G
Cy3-collI uptake
% Cy3-positive
(corrected to running average)
5
0
-5
10
10
20
30
40
50
60
Time (h, post-synchronization)

**Figure S2**

A. iTTFs were incubated with Cy3-colI (magenta) for one hour, trypsinized, replated and allowed to stick down for 18hrs, before being fixed and stained with collagen-I (ColI, green) and counterstained with DAPI (blue). Arrowhead indicating co-localization of labelled collagen-I with endogenous collagen-I. Scale bar = 10 µm. Representative of N=3.

B. Representative images from flow imaging of iTTFs either incubated with Cy5-colI only (top) or both Alexa fluor 488-labelled 70kDa dextran and Cy5-colI (bottom), showing very little co-localization between the two markers after being taken up into cells. Scale bar = 10 µm. N=3, >800 cells analyzed per experiment.

C. Representative images from Flow imaging of iTTFs either incubated with Cy5-colI only (top) or Cy5-colI and unlabelled colI (bottom), showing that in presence of excess unlabelled collagen, the majority of the Cy5 signal are restricted to the periphery of the cells. Scale bar = 10 µm. N=3, >800 cells analyzed per experiment.

D. Scatter plot showing Dyngo4a (Dyng), an endocytosis inhibitor, treatment for one hour inhibits over 50% of Cy3-colI uptake in iTTFs. Bars show mean ± s.e.m. of N = 6. ****$p$<0.0001.

E. Alamar Blue assay showing that prolonged treatment of 20 µM Dyngo4a (Dyng) does not inhibit iTTF proliferation.

F. qPCR analysis of *Col1a1* and *Fn1* mRNA levels in DMSO and Dyng-treated iTTFs, corrected to DMSO control, showing a decrease in both collagen-I and fibronectin transcripts. 2-way ANOVA was carried out. Bars showing mean± s.e.m. of N=3, ****$P<0.0001$.

G. Percentage Cy3-colI taken up by synchronized iTTFs over 48 h. Fluctuation of Cy3-positive cells was corrected to running average (average of 12 h). Bars show mean ± s.e.m. of N = 3 per time point.

Figure S3

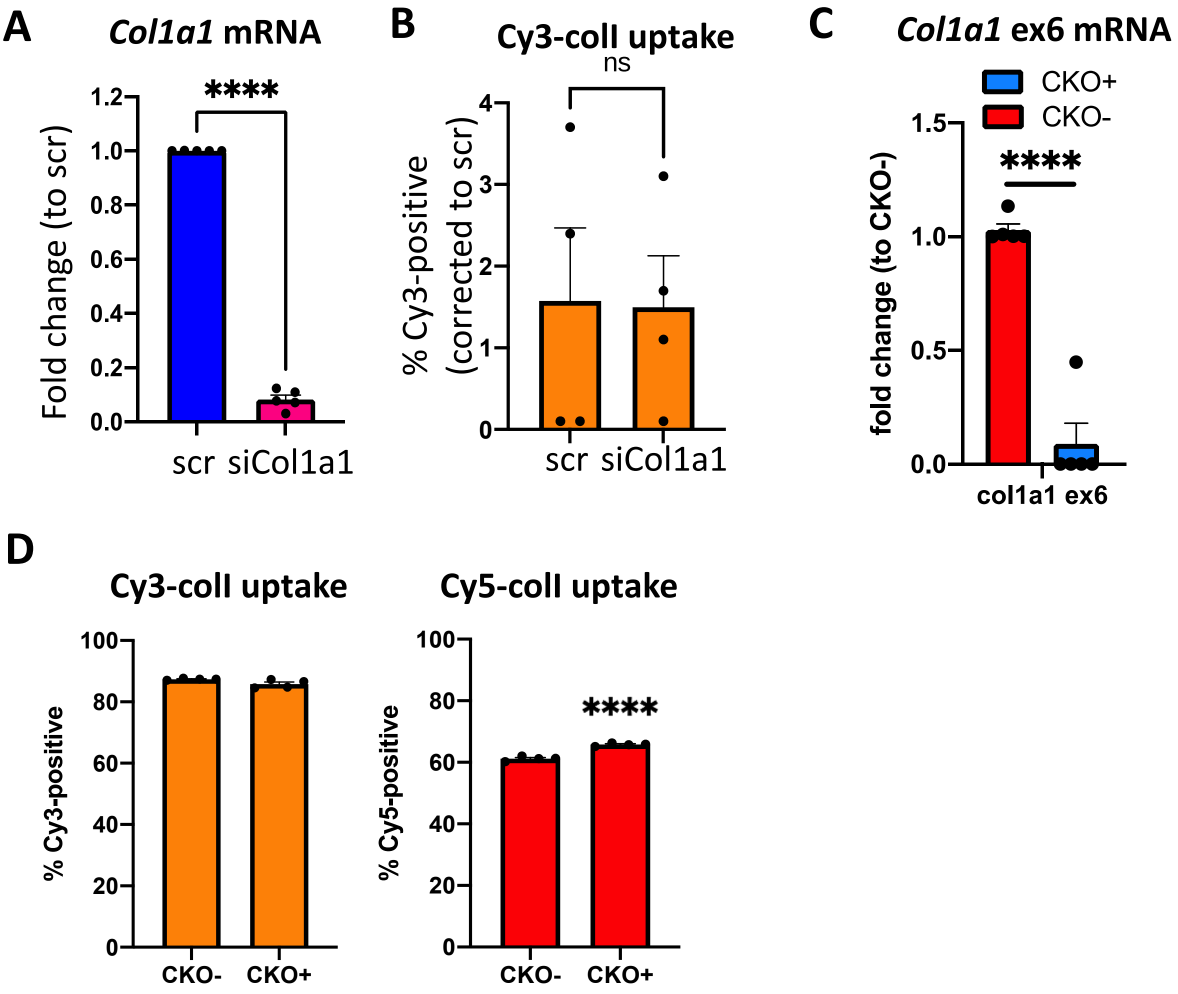
A
Col1a1 mRNA
****
Fold change (to scr)
1.2
1.0
0.8
0.6
0.4
0.2
0.0
scr
siCol1a1
B
Cy3-colI uptake
ns
% Cy3-positive (corrected to scr)
4
3
2
1
0
scr
siCol1a1
C
Col1a1 ex6 mRNA
CKO+
CKO-
****
fold change (to CKO-)
1.5
1.0
0.5
0.0
col1a1 ex6
D
Cy3-colI uptake
% Cy3-positive
100
80
60
40
20
0
CKO-
CKO+
Cy5-colI uptake
% Cy5-positive
****
100
80
60
40
20
0
CKO-
CKO+

**Figure S3**

A. qPCR analysis of scr and siCol1a1 iTTFs. N = 5. $p<0.0001$

B. Bar chart showing the percentage of iTTFs that have taken up 5 µg/mL Cy3-colI (left) or 5 µg/mL Cy5-colI (right) after 1 h incubation. Paired t-test was performed, N = 4. *$p = 0.0372$.

C. qPCR analysis of CKO- and CKO+ primary tail tendon fibroblasts. Col1a1ex6 are primers specific to the locus being knocked out, and col1a1 are primers detecting general collagen-I mRNA. N=5. ****$p<0.0001$.

D. Bar chart showing the % of primary tail tendon fibroblasts that have taken up 5ug/ml Cy3-colI (left) and Cy5-colI (right) after 1 hour incubation; CKO+ cells have a similar uptake to CKO- when incubated with Cy3-colI, and a slight but significant increase in uptake of Cy5-colI. N = 4. ****$p<0.0001$.

Figure S4

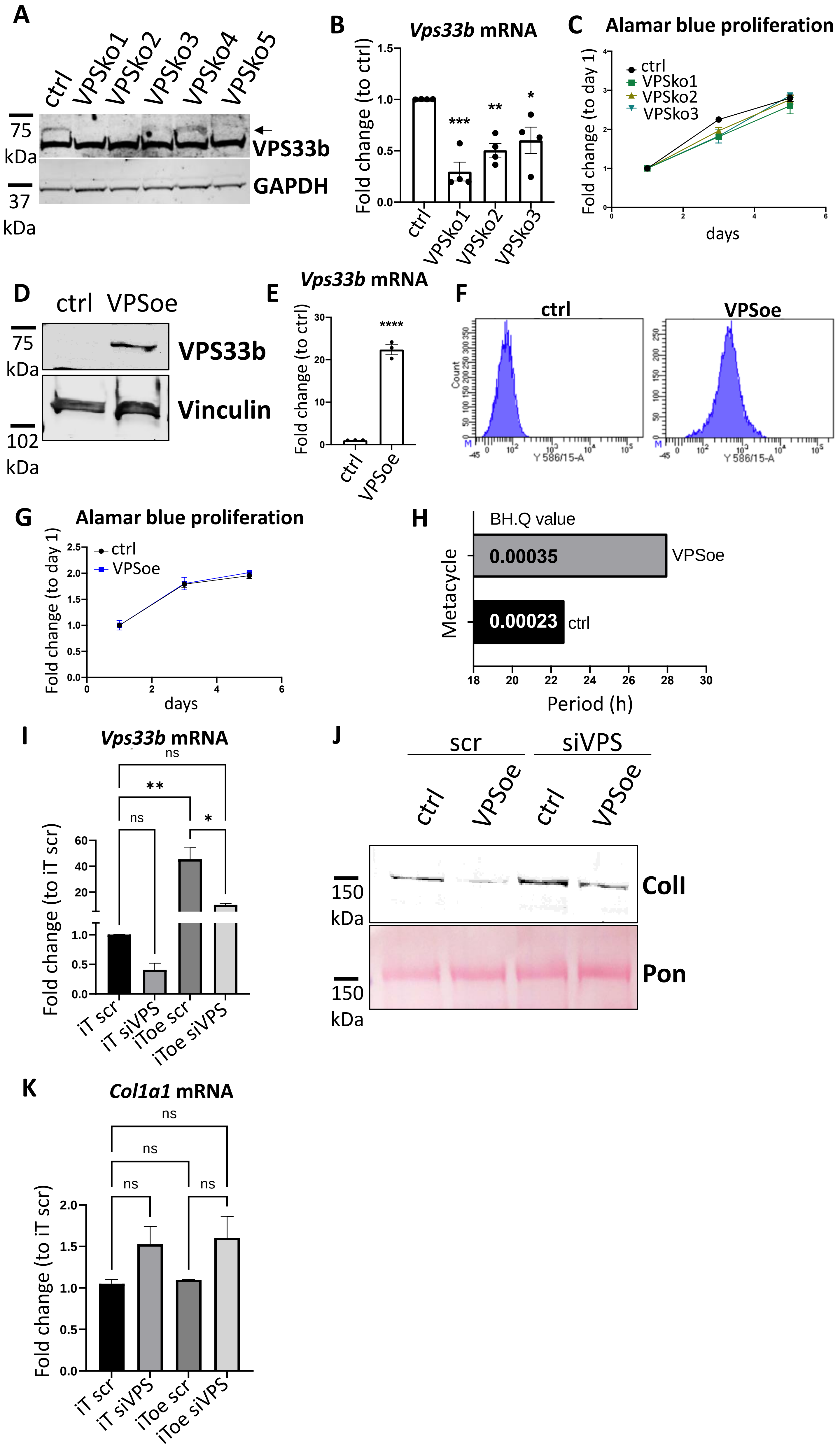

A
ctrl
VPSko1
VPSko2
VPSko3
VPSko4
VPSko5
75 kDa
37 kDa
VPS33b
GAPDH
B
Vps33b mRNA
Fold change (to ctrl)
***
**
*
ctrl
VPSko1
VPSko2
VPSko3
C
Alamar blue proliferation
Fold change (to day 1)
ctrl
VPSko1
VPSko2
VPSko3
days
D
ctrl
VPSoe
75 kDa
102 kDa
VPS33b
Vinculin
E
Vps33b mRNA
Fold change (to ctrl)
****
ctrl
VPSoe
F
ctrl
VPSoe
Count
Y 586/15-A
G
Alamar blue proliferation
Fold change (to day 1)
ctrl
VPSoe
days
H
BH.Q value
0.00035
VPSoe
0.00023
ctrl
Metacycle
Period (h)
I
Vps33b mRNA
ns
**
ns
*
Fold change (to iT scr)
iT scr
iT siVPS
iToe scr
iToe siVPS
J
scr
siVPS
ctrl
VPSoe
ctrl
VPSoe
150 kDa
150 kDa
ColI
Pon
K
Col1a1 mRNA
ns
ns
ns
ns
Fold change (to iT scr)
iT scr
iT siVPS
iToe scr
iToe siVPS

**Figure S4**

A. Western blot analysis of VPS33B knockout (VPSko) clones compared to control (ctrl) iTTFs. Top: probed with VPS33B antibody, bottom: probed with GAPDH antibody. Protein molecular weight ladders to the left (in kDa). Representative of N=3.

B. qPCR analysis of *VPS33B* expression in the 3 selected clones. *** $p$=0.0002, ** $p$=0.0039, * $p$=0.0163

C. Alamar blue analysis of proliferation rates of ctrl and VPSko iTTFs. Representative of N=3.

D. Western blot analysis of VPS33B protein levels in control (ctrl) and VPS33B over-expressing (VPSoe) iTTFs. Top panel probed with VPS33B antibody. Bottom panel re-probed with vinculin antibody. Protein molecular weight ladders to the left (in kDa). Representative of N=4.

E. qPCR analysis of *VPS33B* expression in ctrl and VPSoe iTTFs. N=4, **** $p$<0.0001.

F. Single parameter histograms of Flow cytometry analysis on ctrl (left) and VPSoe (right) iTTFs, showing a shift in increase of RFP fluorescence and thus expression of VPSoe vector. Representative of N>4.

G. Alamar blue analysis of proliferation rates of ctrl iTTFs and iTTF VPSoe. Representative of N=3.

H. MetaCycle analyses of the fibril counts showed a rhythmicity of circa 23 h in ctrl iTTFs compared with circa 28 h in iTTF VPSoe.

I. qPCR analysis of *VPS33b* mRNA expression levels in iTTF or iTTF VPSoe, treated with siRNA scrambled control (iT scr, iToe scr) or siRNA against VPS33b (iT siVPS, iToe siVPS) and cultured for 72 h. N=2.

J. Western blot analysis of conditioned media taken from ctrl and VPSoe iTTFs, treated either with siRNA scrambled control (scr) or siRNA against VPS33B (siVPS) and cultured for 72 h. Top: probed with collage-I antibody (ColI), bottom: counterstained with Ponceau (Pon) as control. Protein molecular weight ladders to the left (in kDa). Representative of N=2.

K. qPCR analysis of *Col1a1* mRNA expression levels in iTTF or iTTF VPSoe, treated with siRNA scrambled control (iT scr, iToe scr) or siRNA against VPS33b (iT siVPS, iToe siVPS) and cultured for 72 h. N=2.

### Figure S5

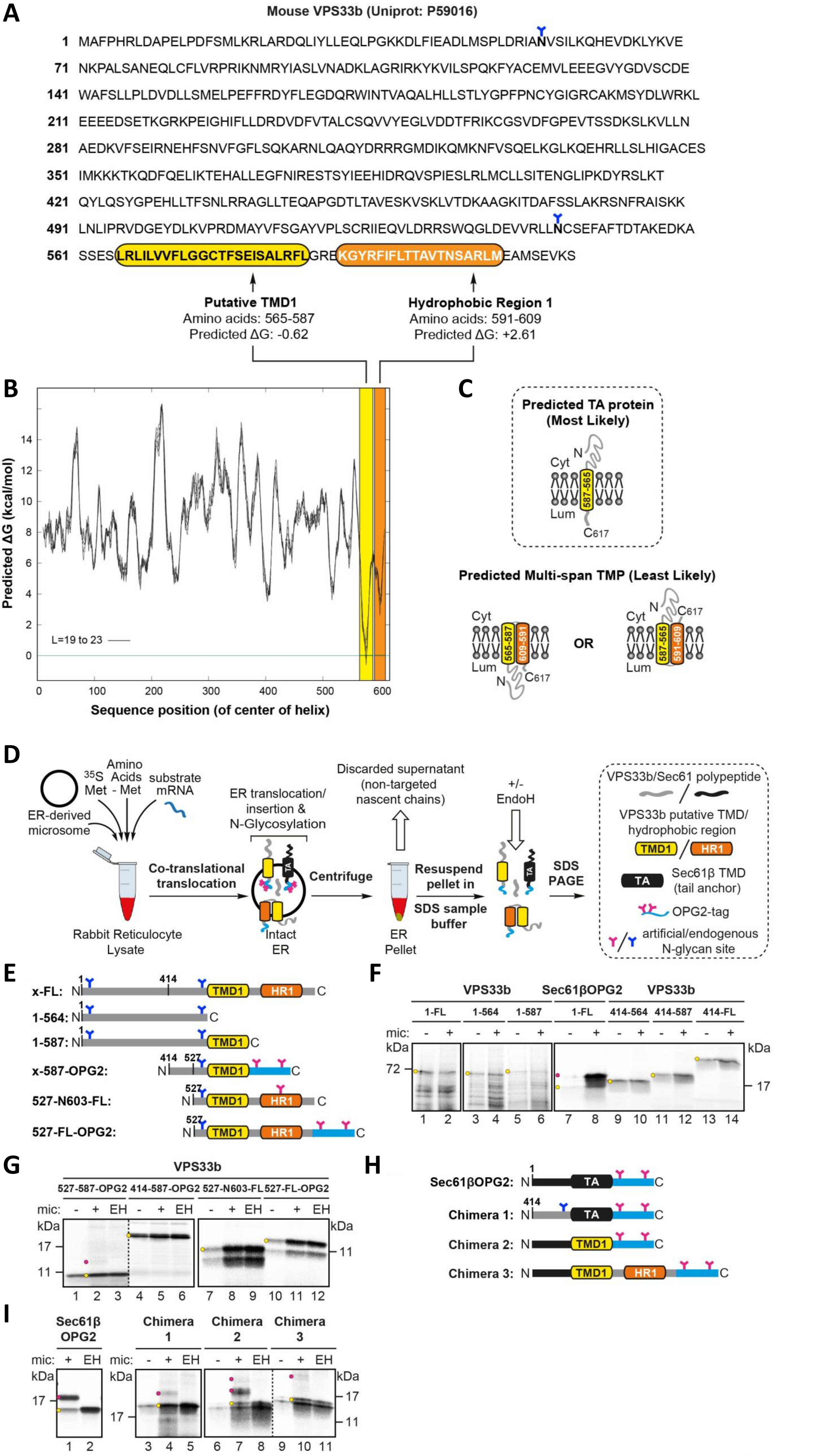
A
Mouse VPS33b (Uniprot: P59016)
1 MAFPHRLDAPELPDFSMLKRLARDQLIYLLEQLPGKKDLFIEADLMSPLDRIANVSILKQHEVDKLYKVE
71 NKPALSANEQLCFLVRPRIKNMRYIASLVNADKLAGRIRKYKVILSPQKFYACEMVLEEEGVYGDVSCDE
141 WAFSLLPLDVDLLSMELPEFFRDYFLEGDQRWINTVAQALHLLSTLYGPFPNCYGIGRCAKMSYDLWRKL
211 EEEEDSETKGRKPEIGHIFLLDRDVDFVTALCSQVVYEGLVDDTFRIKCGSVDFGPEVTSSDKSLKVLLN
281 AEDKVFSEIRNEHFSNVFGFLSQKARNLQAQYDRRRGMDIKQMKNFVSQELKGLKQEHRLLSLHIGACES
351 IMKKKTKQDFQELIKTEHALLEGFNIRESTSYIEEHIDRQVSPIESLRLMCLLSITENGLIPKDYRSLKT
421 QYLQSYGPEHLLTFSNLRRAGLLTEQAPGDTLTAVESKVSKLVTDKAAGKITDAFSSLAKRSNFRAISKK
491 LNLIPRVDGEYDLKVPRDMAYVFSGAYVPLSCRIIEQVLDRRSWQGLDEVVRLLNCSEFAFTDTAKEDKA
561 SSESLRLILVVFLGGCTFSEISALRFLGREKGYRFIFLTTAVTNSARLMEAMSEVKS
Putative TMD1
Amino acids: 565-587
Predicted ΔG: -0.62
Hydrophobic Region 1
Amino acids: 591-609
Predicted ΔG: +2.61
B
Predicted ΔG (kcal/mol)
L=19 to 23
Sequence position (of center of helix)
C
Predicted TA protein (Most Likely)
Cyt
Lum
N
C617
587-565
Predicted Multi-span TMP (Least Likely)
565-587
609-591
OR
587-565
591-609
D
ER-derived microsome
35S Met
Amino Acids - Met
substrate mRNA
Rabbit Reticulocyte Lysate
Co-translational translocation
ER translocation/ insertion & N-Glycosylation
Intact ER
Centrifuge
Discarded supernatant (non-targeted nascent chains)
ER Pellet
Resuspend pellet in SDS sample buffer
+/- EndoH
SDS PAGE
VPS33b/Sec61 polypeptide
VPS33b putative TMD/ hydrophobic region
TMD1
HR1
Sec61β TMD (tail anchor)
TA
OPG2-tag
artificial/endogenous N-glycan site
E
x-FL:
1-564:
1-587:
x-587-OPG2:
527-N603-FL:
527-FL-OPG2:
F
VPS33b
Sec61βOPG2
1-FL
1-564
1-587
414-564
414-587
414-FL
mic:
kDa
72
17
G
VPS33b
527-587-OPG2
414-587-OPG2
527-N603-FL
527-FL-OPG2
EH
11
H
Sec61βOPG2:
Chimera 1:
Chimera 2:
Chimera 3:
I
Sec61β OPG2
Chimera 1
Chimera 2
Chimera 3

**Figure S5**

A. The amino acid sequence of mouse VPS33B as listed in Uniprot (P59016). Each predicted putative transmembrane domain (TMD) is highlighted in yellow (TMD1) and an adjacent hydrophobic region in orange (HR2), together with their respective predicted ΔG values (kcal/mol) for TMD insertion in the endoplasmic reticulum (ER) membrane (see part B). Potential consensus sites for the addition of N-linked glycans (N-X-S/T) are denoted by a blue Y symbol.

B. The amino acid sequence in A was subjected to the full scan option of the ΔG prediction server for identifying potential transmembrane helices: http://dgpred.cbr.su.se/.

C. Potential topologies of mouse VPS33B in the ER membrane. Based on the predicted ΔG values estimated in B, HR1 is predicted to be poorly inserted into the ER membrane. Hence, VPS33B most likely acquires a tail-anchor (TA) protein topology (see hashed box) as opposed to that of a multi-span transmembrane protein (TMP). Cyt, cytosol; Lum, ER lumen.

D. Outline of the *in vitro* assay using canine pancreatic as a source of ER membrane; following translation, membrane inserted radiolabelled precursor proteins are recovered by centrifugation and analyzed by SDS-PAGE and phosphorimaging. The N-glycosylation of lumenal domains, confirmed by treatment with endoglycosidase H (Endo H), indicates successful membrane translocation.

E. Schematics of endogenous, truncated and OPG2 tagged VPS33b proteins used in this study.

F. Non-glycosylated and N-glycosylated radiolabelled cell free translation products are respectively indicated by a yellow or magenta circle. The substrates depicted in E, and the model tail-anchor (TA) protein Sec61β (modified with a C-terminal OPG2 tag), were synthesized as outlined in D in the absence (odd lanes) and presence (even lanes) of canine pancreatic microsomes (indicated as – or + mic). In each case, a significant proportion of full-length and truncated forms of VPS33b pelleted in the absence of microsomes (indicative of aggregation).

G. Non-glycosylated and N-glycosylated radiolabelled products are respectively indicated by a yellow or magenta circle. The OPG2 tagged truncated forms of VPS33b depicted in B were synthesized as described in C and treated with EndoH (EH) (lanes 3, 6, 9 and 12). In no case were any domains N-glycosylated.

H. Schematics of OPG2 tagged VPS33b and Sec61β chimeric proteins used in this study; chimera 1: residues 414-564 VPS33b, residues 73-94 Sec61β (TA region), OPG2 tag; chimera 2: residues 1-72 Sec61β (N-terminal region), residues 565-587 VPS33b (TMD1), OPG2 tag; chimera 3: residues 1-72 Sec61β (N-terminal region), residues 565-617 VPS33b (TMD1, HR1 and C-terminus), OPG2 tag.

I. Non-glycosylated and N-glycosylated radiolabelled products are respectively indicated by a yellow or magenta circle. Sec61βOPG2 and chimeras 1-3 depicted in H were synthesized and treated with EndoH (EH) as outlined in D.

#### Figure S6

**A**

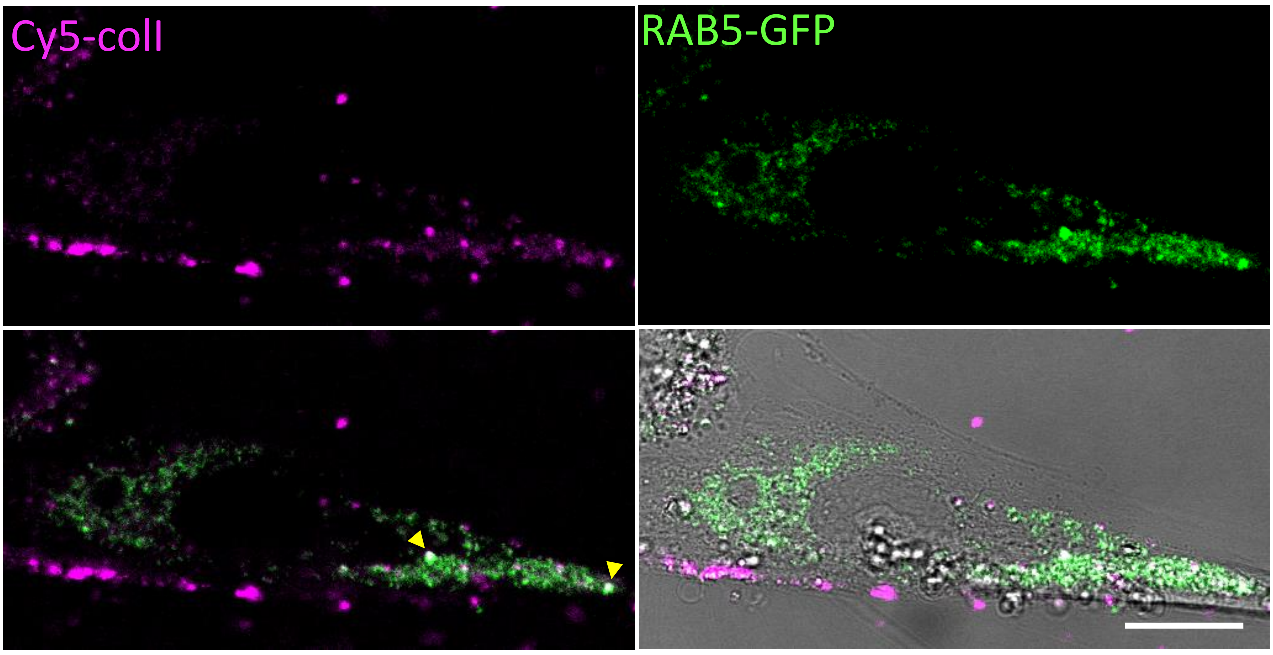

Cy5-coll
RAB5-GFP


**A**

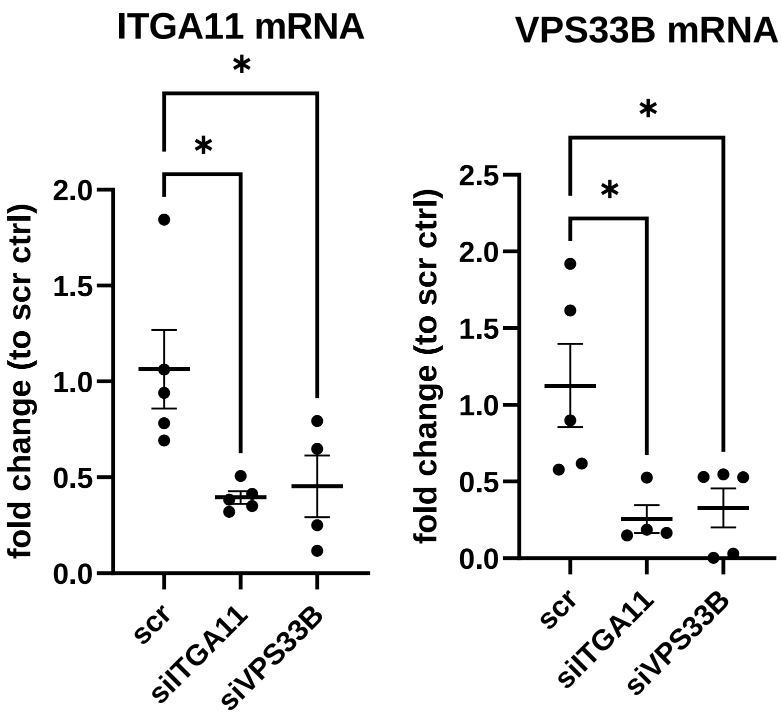

ITGA11 mRNA
VPS33B mRNA
fold change (to scr ctrl)
2.0
1.5
1.0
0.5
0.0
2.5
2.0
1.5
1.0
0.5
0.0
*
*
*
*
scr
siITGA11
siVPS33B
scr
siITGA11
siVPS33B

**Figure S6**

A. Human lung fibroblasts transiently transfected with GFP-tagged RAB5 (RAB5-GRP, green) and with Cy5-colI added before confocal live imaging. Individual fluorescence channels were presented here together with merged images on the bottom row (with or without brightfield). Yellow arrowheads point out intracellular structures where Cy5-colI co-localises with RAB5-GFP. Scale bar = 10 µm.

B. qPCR analyses of ITGA11 and VPS33B mRNA levels in IPF cells treated with siRNA against ITGA11 (siITGA11) or siRNA against VPS33B (siVPS33B). Ordinary one-way ANOVA analysis with multiple comparisons performed, n=5 across N=2. *ITGA11*, siITGA11 *$p = 0.0145$, siVPS33B *$p = 0.0319$; *VPS33B*, siITGA11 *$p = 0.018$, siVPS33B *$p = 0.0207$;

### Figure S7

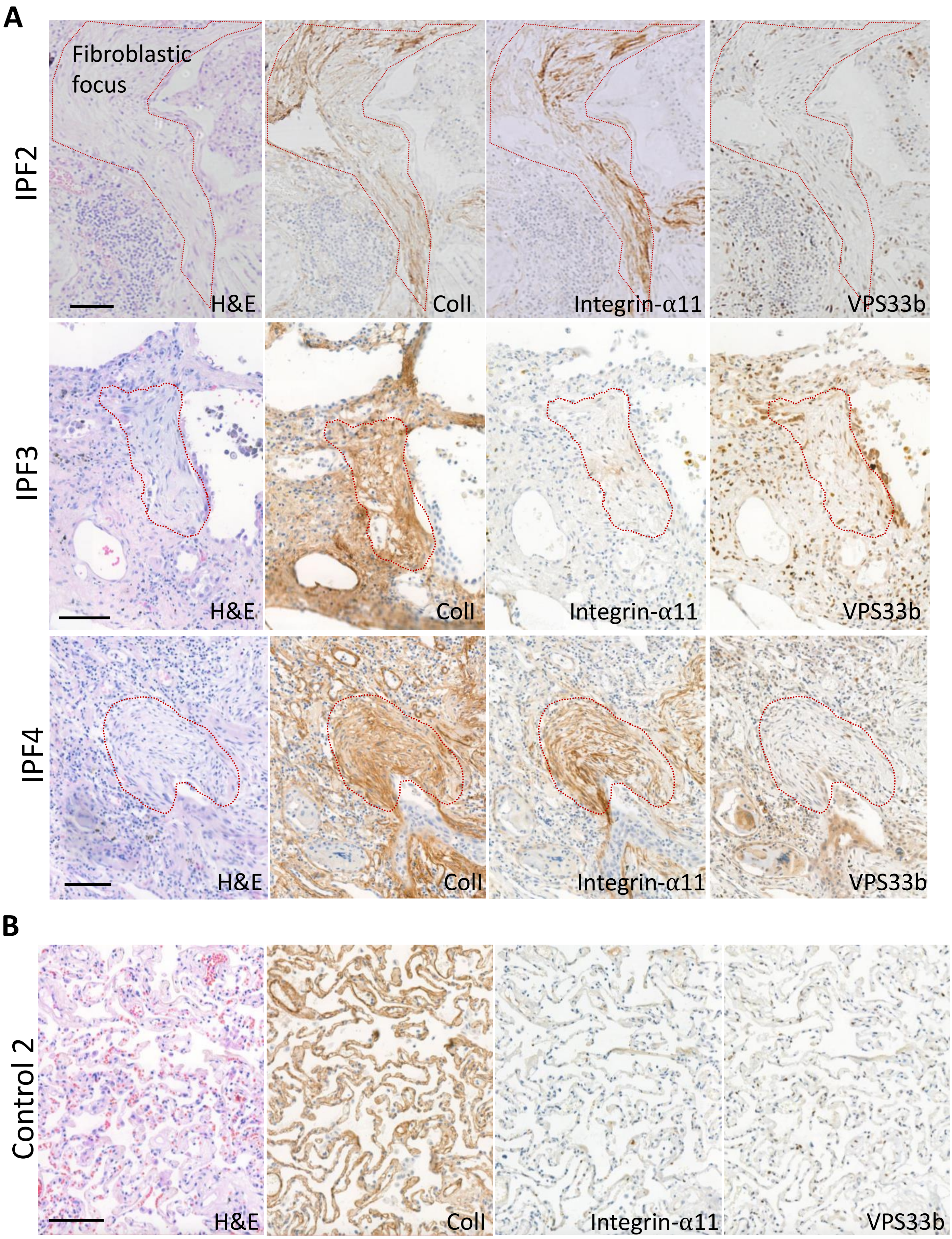
A
IPF2
Fibroblastic focus
H&E
ColI
Integrin-α11
VPS33b
IPF3
H&E
ColI
Integrin-α11
VPS33b
IPF4
H&E
ColI
Integrin-α11
VPS33b
B
Control 2
H&E
ColI
Integrin-α11
VPS33b

**Figure S7**

A. Immunohistochemistry of 5 µm thick sequential lung sections taken from 3 additional IPF patients (IPF2, IPF3, IPF4). Red dotted lines outline the fibroblastic focus. Sections were stained with hematoxylin and eosin (H&E), collagen-I (ColI), integrin-$\alpha$11, VPS33B. Scale bar = 100 µm.

B. Immunohistochemistry of 5 µm thick sequential lung sections taken from an additional lung classified as control (Control 2). Sections were stained with hematoxylin and eosin (H&E), collagen-I (ColI), integrin-$\alpha$11, VPS33B. Scale bar = 100 µm.
